# The RAD51 recombinase protects mitotic chromatin in human cells

**DOI:** 10.1101/2020.08.11.246231

**Authors:** Isabel E. Wassing, Xanita Saayman, Lucia Rampazzo, Christine Ralf, Andrew Bassett, Fumiko Esashi

## Abstract

The RAD51 recombinase plays critical roles in safeguarding genome integrity, which is fundamentally important for all living cells. While interphase functions of RAD51 in repairing broken DNA and protecting stalled replication forks are well characterised, its role in mitosis remains contentious. In this study, we show that RAD51 protects under-replicated DNA in mitotic human cells and, in this way, promotes mitotic DNA synthesis (MiDAS) and successful chromosome segregation. MiDAS was globally detectable irrespective of DNA damage and was promoted by *de novo* RAD51 recruitment, RAD51-mediated fork protection, and RAD51 phosphorylation by the key mitotic regulator Polo-like kinase 1. Importantly, acute inhibition of RAD51-promoted MiDAS led to mitotic DNA damage, delayed anaphase onset and induced centromere fragility, revealing a mechanism that prevents the satisfaction of the spindle assembly checkpoint when chromosomal replication remains incomplete. This study hence identifies an unexpected function of RAD51 in promoting the stability of mitotic chromatin.

## Introduction

The RAD51 recombinase is essential for cell survival and for the protection of genome stability. RAD51 is best known as the central catalyst of homologous recombination (HR), which provides error-free repair of double-stranded DNA breaks (DSBs). In this process, DSBs are first recognised by the MRN complex, comprising MRE11, RAD50 and NBS1, which, in cooperation with CtIP, resects the broken DNA ends during S and G2 phases of the cell cycle (Ceccaldi et al., 2016; Garcia et al., 2011; Shibata et al., 2014). The resected 3’ single-stranded DNA (ssDNA) overhang is rapidly bound by replication protein A (RPA), which is subsequently replaced by RAD51. The resulting RAD51 nucleoprotein filament then mediates strand invasion to initiate HR (Baumann and West, 1998). This mechanism enables RAD51 to catalyse HR not only at two-ended DSBs, but also at one-ended DSBs arising from collapsed replication forks, thereby promoting replication restart through break-induced replication (BIR) (Benedict et al., 2020; Hashimoto et al., 2011; Natsume et al., 2017). RAD51 also displays non-repair functions at stalled replication forks, in which RAD51 stimulates fork reversal and stabilises stalled forks by protecting against the nucleolytic degradation of nascent DNA (Mason et al., 2019; Morrison et al., 1999; Petermann et al., 2010; Schlacher et al., 2011; Schlacher et al., 2012; Wang et al., 2015).

In interphase, RAD51 loading onto RPA-coated ssDNA is largely mediated by the breast cancer susceptibility 2 protein (BRCA2) (Davies and Pellegrini, 2007; Esashi et al., 2007; Jensen et al., 2010), which is recruited to sites of DSB through its interaction with the partner and localizer of BRCA2 (PALB2) and the breast cancer susceptibility 1 protein (BRCA1) (Zelensky et al., 2014). As cells progress into mitosis, however, RAD51 binding to BRCA2 is perturbed by CDK-mediated BRCA2 phosphorylation, suggesting that BRCA2-mediated RAD51 loading to chromatin is downregulated in mitosis (Esashi et al., 2005). These observations are in agreement with the reported attenuation of the DNA damage response (DDR) in mitosis; recruitment of DSB repair mediators such as BRCA1 or 53BP1, which promote HR or error-prone non-homologous end joining (NHEJ), respectively, is perturbed in mitotic cells. Interestingly, restoration of 53BP1 recruitment to mitotic chromatin results in telomere fusions and reduced cell survival following irradiation, indicating that the activation of NHEJ in mitosis is deleterious (Benada et al., 2015; Lee et al., 2014; Orthwein et al., 2014). However, it remains unclear whether the recruitment of HR proteins in mitosis is similarly disadvantageous.

Significantly, our previous studies uncovered a mechanism that enables RAD51 recruitment to damaged DNA independently of BRCA2 (Yata et al., 2014; Yata et al., 2012). This pathway is triggered by a key mitotic regulator, Polo-like kinase 1 (PLK1), which phosphorylates RAD51 at serine 14 (S14) shortly after DNA damage and in later phases of the cell cycle. This event provokes RAD51 phosphorylation by the acidophilic casein kinase 2 (CK2) at threonine 13 (T13), which mediates a direct interaction between RAD51 and the NBS1 component of the MRN complex. In this way, PLK1-mediated RAD51 phosphorylation allows RAD51 to be recruited to broken DNA ends and also to stalled replication forks (Yata et al., 2014). Given that RAD51 phosphorylation peaks during late G2 phase and mitosis in unperturbed cells (Yata et al., 2014; Yata et al., 2012), these observations raise the intriguing possibility that, similar to the well-described role of RAD51 in meiosis (Dai et al., 2017), RAD51-mediated break repair may be enabled in mitosis.

In recent years, there has been an increasing appreciation that, upon exposure to mild replicative stress, cancer cells enter mitosis before the completion of DNA replication (Bergoglio et al., 2013; Bhat et al., 2013; Naim et al., 2013). The persistence of under-replicated DNA in mitosis, often described at common fragile sites (CFSs), telomeres and centromeres, leads to impaired chromosome disjunction, observed as anaphase bridges, ultra-fine bridges (UFBs) and micronuclei (Bergoglio et al., 2013; Lukas et al., 2011; Minocherhomji et al., 2015; Naim and Rosselli, 2009; Naim et al., 2013). Several proteins have been found to act at CFSs to alleviate these phenotypes, such as FANCD2, MUS81, TOPBP1 and TLS polymerases Pol η (eta) and Pol ζ (zeta) (Bergoglio et al., 2013; Bhat et al., 2013; Chan et al., 2009; Naim and Rosselli, 2009; Pedersen et al., 2015; Ying et al., 2013). It has been proposed that these factors work in concert at under-replicated CFSs to complete replication and/or promote mitotic DNA synthesis (MiDAS) via a BIR-like mechanism, triggered by nucleolytic cleavage at stalled replication forks (Dilley et al., 2016; Min et al., 2017; Minocherhomji et al., 2015). While several reports in higher eukaryotes suggest that RAD51 catalyses BIR, MiDAS at CFSs and telomeres appears to be primarily dependent on RAD52 to complete replication in early mitosis (Benedict et al., 2020; Bhowmick et al., 2016; Costantino et al., 2014; Min et al., 2017; Natsume et al., 2017).

Given that cancer cells generally experience increased levels of oncogene-induced replication stress, MiDAS is thought to be particularly important for their survival, making the inhibition of MiDAS an attractive avenue for cancer therapy (Bhowmick et al., 2016). Nonetheless, a complete picture of the mechanisms orchestrating MiDAS is currently lacking. In this study, we show that RAD51 promotes MiDAS at both broken and unbroken replication forks, and that this involves RAD51-mediated fork protection. Analysis of RAD51 missense mutants at S14 further indicates that RAD51 phosphorylation by PLK1 mediates this mitotic RAD51 function. Additionally, we find that inhibition of the spindle assembly checkpoint (SAC) severely decreases cell survival in cells that exhibit high levels of replication stress and that mitotic under-replication impacts mitotic progression. Lastly, the conditions that inhibit MiDAS induce fragility at centromeres, providing a potential link between replication completion and SAC maintenance. On the basis of these findings, we propose that RAD51 plays an active role in mitosis by protecting under-replicated DNA from nucleolytic attack. In this way, RAD51 promotes MiDAS to ensure completion of replication at these loci, thereby regulating mitotic progression and facilitating accurate chromosome inheritance.

## Results

### MiDAS in U2OS cells is largely associated with unbroken DNA

We first aimed to determine the nature of MiDAS in further detail. To this end, U2OS cells were synchronised at G1/S by double thymidine block and then released into S phase (dT-B/R) in medium containing a low dose (0.4 µM) of aphidicolin (APH), an inhibitor of B-family DNA polymerases α, δ and ε. The CDK1 inhibitor RO3306 was then added as the cells entered G2 phase to arrest them at the G2/M boundary, followed by mitotic release (m-R) into fresh medium containing the thymidine analog EdU. This procedure allowed robust enrichment of mitotic cells (**Figure S1A**). Cells were then collected by mitotic shake off (m-SO) and EdU incorporation was analysed by fluorescent click-labelling (EdU-Click-F). To assess whether the EdU foci observed represent DNA synthesis in late G2 or mitosis, the EdU pulse was administered in G2 phase (30 min prior to m-R) or upon m-R for varying lengths of time (**Figure S1B-C**). While a substantial amount of EdU signal was observed in cells pulse labelled in G2, this signal was drastically reduced when the G2-pulse was followed by a thymidine chase upon m-R, suggesting that little EdU incorporation occurs during the RO3306 block. Conversely, strong EdU signals were detectable when cells were exposed to EdU concurrently with m-R, while these levels were proportionally decreased when EdU addition was delayed for 5 min or longer. These observations suggest that MiDAS predominantly occurs during the very early stages of mitosis.

We next tested whether the duration of RO3306 incubation affected MiDAS (**Figure S1D-E**). Similar to previous reports (Garribba et al., 2018), a significant reduction of mitotic EdU incorporation was detectable when cells were arrested by RO3306 for longer (**Figure S1E**), supporting the view that APH-induced under-replication can, at least in part, be resolved by extending the duration of G2. Mitotic *γ*-H2AX, an early and sensitive marker for the DDR, was similarly reduced upon prolonged RO3306-arrest (**Figure S1F**). Nonetheless, a substantial amount of mitotic EdU incorporation remained even after 12 hours of RO3306 incubation, suggesting that a subset of under-replicated regions necessarily require release from RO3306, or mitotic entry, to be fully resolved. Intriguingly, we noticed that, for all the different synchronisation protocols tested for U2OS cells, only a subset of EdU foci (< 30%) colocalised with *γ*-H2AX (**Figure S1G**). In HEK293 cells, however, the majority of mitotic EdU foci (64%) were *γ*-H2AX-positive (**Figure S1H**). These observations suggest that the nature of MiDAS varies significantly between different cell lines; MiDAS is largely associated with unbroken DNA in U2OS, but not in HEK293.

### RAD51 promotes MiDAS

We next set out to investigate the potential involvement of RAD51 in MiDAS. To this end, U2OS cells were synchronised in combination with siRNA treatment such that mitotic cells were collected 48 hours after transfection (**Figure 1A**). As prolonged RAD51 depletion was expected to trigger G2 arrest due to its essential role during DNA replication (Sonoda et al., 1998; Tsuzuki et al., 1996), cells were also treated with the WEE1 inhibitor AZD1775 (previously known as MK-1775) to bypass this arrest. Western blot analysis confirmed the successful depletion of RAD51 from the total cell population (**Figure 1B**). As expected, cells treated with siRNA targeting RAD51 (siR51) showed a reduced mitotic population compared to cells treated with a negative control siRNA (siCntl), as detected by histone H3 phosphorylation at serine 10 (pS10-H3), and this impact was alleviated by WEE1 inhibition (**Figure 1B and C**). We found that mitotic EdU incorporation was indeed significantly decreased in siR51-treated cells compared to control cells (**Figure 1D and E**). In contrast, mitotic *γ*-H2AX foci appeared unchanged (**Figure 1D and F**) and mitotic RPA foci were increased (**Figure 1D and G**). The same trend was observed in cells co-treated with WEE1 inhibitor, with the exception of mitotic *γ*-H2AX foci, which showed a modest decrease in siR51-treated cells (**Figure 1F**). While these data suggest that RAD51 depletion led to defects in both completing DNA replication and MiDAS, we were unable to exclude the possibility that the delayed progression of S and G2 phase under these conditions prevents mitotic entry of severely affected cells, even in the presence of WEE1 inhibitor, such that m-SO would result in the collection of an unrepresentative sub-population in the siR51-treated samples compared to the corresponding control. This may explain the decrease in mitotic *γ*-H2AX foci observed in RAD51-depleted cells (**Figure 1F and S2G**).

**Figure 1:**
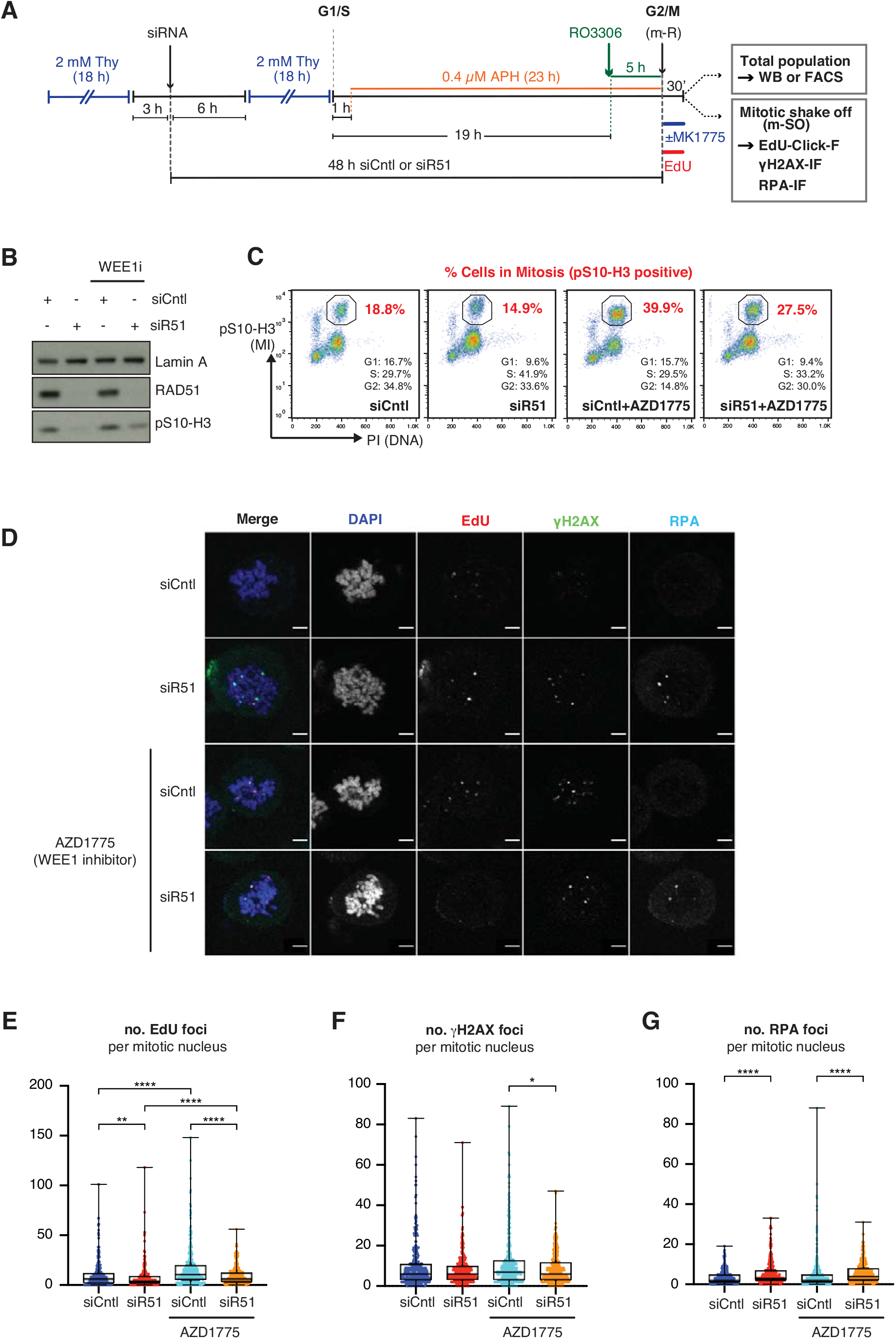
RAD51 depletion significantly reduces MiDAS. **A.** Schematic diagram of U2OS cell synchronisation by double thymidine block and release (dT-B/R) with siRNA treatment against RAD51 (siR51) or control (siCntl), followed by exposure to low-dose aphidicolin (0.4 µM APH) and G2/M arrest by CDK1 inhibitor (9 µM, RO3306). Cells were released into mitosis (m-R) in medium containing 10 µM EdU with or without 100 nM WEE1 inhibitor (100 nM, AZD1775). **B-C**. Western blot analysis against RAD51, mitotic marker pS10-H3 and Histone H3 (B) or FACS analysis for mitotic index (pS10-H3) and DNA content (PI) (C) of the total cell population 30 min after m-R, according to the workflow depicted in (A). **D.** Representative images of cells collected by mitotic shake-off (m-SO) at 30 min after m-R, stained for EdU incorporation and *γ*-H2AX and RPA. Scale bar indicates 5 µm. **E-G.** Quantification of the number of EdU foci (E), *γ*-H2AX foci (F) and RPA foci (G) per mitotic nucleus. Data were obtained from three independent experiments (100 cells analysed per repeat); n=300 per condition. Data distribution is represented by box-and-whisker plots (whiskers spanning min-max range of values). Mann Whitney test p-values values are shown. Asterisks indicate p value ≤ 0.05 = *; ≤ 0.01 = **; ≤ 0.001 = ***; ≤ 0.0001 = ****.

In order to circumvent the potential bias in mitotic cells collected upon prolonged depletion of RAD51, we next exploited a well-characterised small molecule inhibitor of RAD51, B02 (Huang et al., 2011) (**Figure 2A**). We reasoned that the use of this small molecule inhibitor would allow for the rapid inhibition of RAD51 during mitosis, without affecting RAD51 functions during the previous S phase. Previous biochemical analyses have shown that B02 inhibits both RAD51 binding to ssDNA and the RAD51-driven strand invasion process, but does not disrupt an already established RAD51 nucleoprotein filament (Huang et al., 2012). In line with these reported properties of B02, damage-induced RAD51 foci formation was significantly hindered when B02 was added prior to, but not after, irradiation (**Figure S2A-C**). As depicted in **Figure 2B**, B02 therefore specifically inhibits *de novo* RAD51 recruitment to DNA, without impacting existing RAD51 filaments. Importantly, we found that addition of B02 upon m-R drastically reduced mitotic EdU incorporation, indicating that *de novo* RAD51 recruitment to mitotic chromatin promotes MiDAS (**Figure 2C**). A corresponding increase in *γ*-H2AX foci was observed upon mitotic RAD51 inhibition, indicating the importance of *de novo* RAD51 recruitment in preventing the accumulation of DNA damage during mitosis (**Figure 2D**).

**Figure 2:**
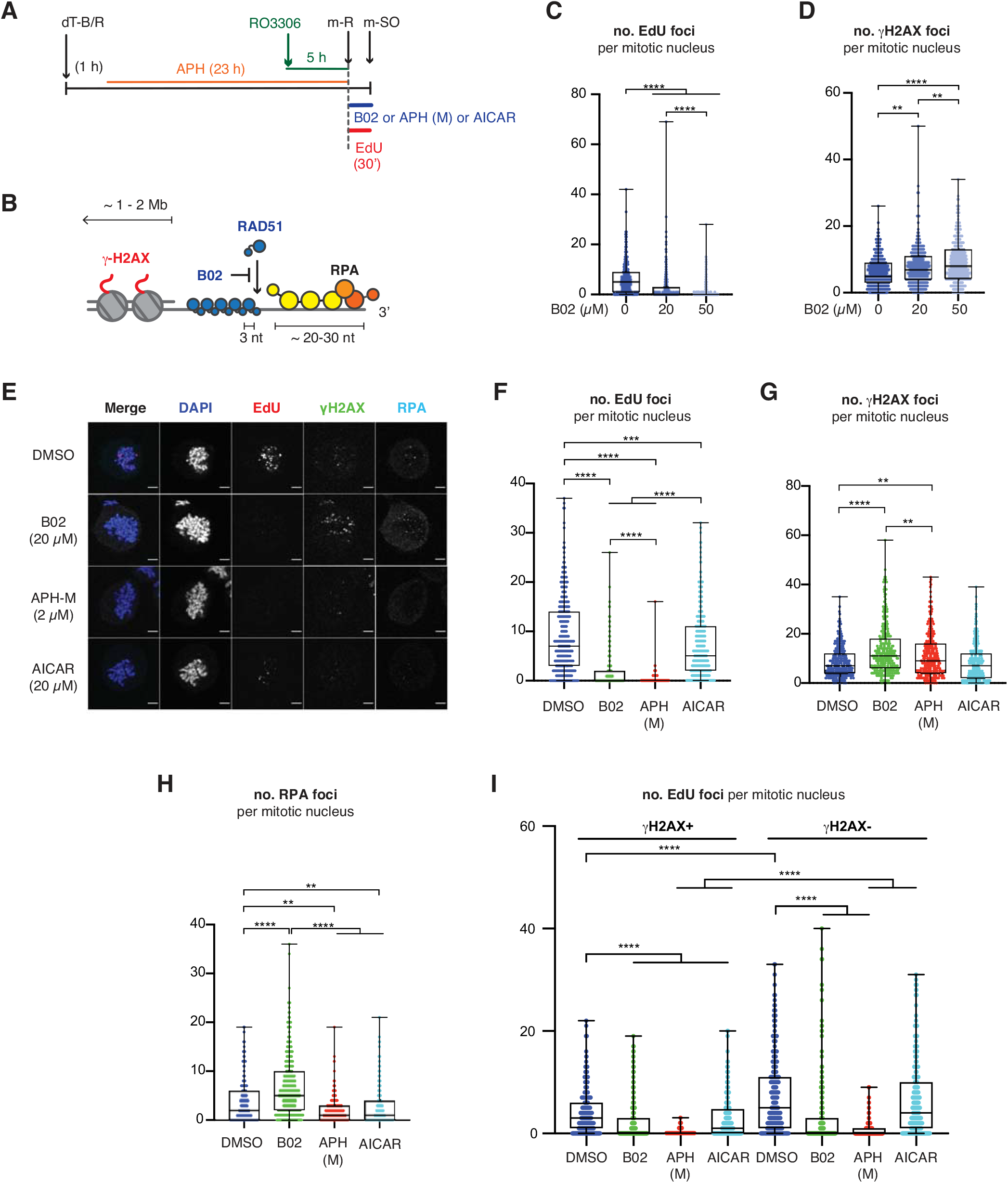
Break-associated and non-break-associated MiDAS are RAD51-dependent. **A.** Schematic diagram of experimental procedure, as in Figure 1A, except, instead of siRNA treatments, cells were exposed to small molecule inhibitors of RAD51 (B02), B-family polymerases (2 µM APH (M)) or RAD52 (20 µM AICAR) upon m-SO. **B.** Depiction of RAD51 and RPA association with resected ssDNA with indicated footprints, and associated *γ*-H2AX, spanning an estimated 1-2 Mb. B02 blocks *de novo* RAD51 loading, but does not disrupt pre-formed filament. **C-D.** Mitotic cells collected by m-SO, following 10 µM EdU exposure with or without B02 at the indicated concentrations, were stained and quantified for EdU foci (C) or *γ*-H2AX foci (D) per mitotic nucleus. The G1, S and G2 populations were estimated by the Watson Pragmatic algorithm (FlowJo), based on the PI staining of interphase cells. **E**. Representative images of mitotic cells collected by m-SO, following 10 µM EdU exposure with or without indicated small molecule inhibitors, stained for EdU incorporation and *γ*-H2AX and RPA. Scale bar indicates 5 µm. **F-I.** Quantification of the number of EdU foci (F), *γ*-H2AX foci (G) and RPA foci (H) per mitotic nucleus. The same data as in (F) and (G) is quantified in (I) as the number of EdU foci that colocalise with *γ*-H2AX (*γ*-H2AX^+^ EdU foci) and the number of EdU foci that do not colocalise with *γ*-H2AX (*γ*-H2AX^-^ EdU foci) per mitotic nucleus. Data were obtained from three independent experiments (100 cells analysed per repeat); n=300 per condition. Data distribution is represented by box-and-whisker plots (whiskers spanning min-max range of values). Mann Whitney test p-values values are shown. Asterisks indicate p value ≤ 0.05 = *; ≤ 0.01 = **; ≤ 0.001 = ***; ≤ 0.0001 = ****.

### RAD52 specifically promotes break-associated MiDAS

MiDAS is widely considered to be a POLD3- and RAD52 -dependent process (Bhowmick et al., 2016; Minocherhomji et al., 2015) and predominantly occurs at CFSs and telomeres via a BIR-like mechanism (Costantino et al., 2014; Dilley et al., 2016; Min et al., 2017). To better understand the relative contributions of these factors in the context of global MiDAS, we compared the effects of B-family polymerase, RAD51 and RAD52 inhibition. As reported previously, MiDAS was fully abolished upon the addition of high-dose (2 µM) aphidicolin in mitosis (APH-M) (**Figure 2E and F**). Interestingly, APH-M treatment increased mitotic *γ*-H2AX foci (**Figure 2G**), suggesting that the *γ*-H2AX foci observed at sites of mitotic EdU incorporation are unlikely to represent fork collapse during DNA synthesis in mitosis. Instead, these foci likely represent DNA breaks that were induced prior to MiDAS, triggering the BIR-like MiDAS mechanism. Conversely, APH-M treatment decreased the number of RPA foci (**Figure 2H**), suggesting that the mitotic RPA signal largely arises due to ongoing MiDAS events. In contrast, similar to siRNA-mediated RAD51 depletion, the number of RPA foci was significantly increased upon mitotic RAD51 inhibition, likely indicating the lack of RAD51-mediated protection of under-replicated regions against nucleolytic degradation, leading to the accumulation of ssDNA under these conditions.

We noticed that, while both mitotic RAD51 and RAD52 inhibition by small molecule inhibitors i.e., B02 and AICAR, respectively, significantly reduced MiDAS (**Figure 2F**), the impact of RAD52 inhibition was limited to EdU foci colocalising with *γ*-H2AX (**Figure 2I**). A similar trend was observed when RAD52 was depleted by siRNA (**Figure S2D-H**). These observations suggest that RAD52 contributes to a specific break-induced MiDAS mechanism, which constitutes a fraction of all global MiDAS detectable in U2OS cells. Strikingly, RAD51 inhibition significantly reduced both *γ*-H2AX-colocalising and non-*γ*-H2AX-colocalising EdU foci, indicating that RAD51 promotes MiDAS in a wider context.

### The role of RAD51 in fork protection promotes MiDAS

The increase in *γ*-H2AX foci upon mitotic RAD51 inhibition prompted us to question whether the importance of RAD51 in MiDAS depends on the recombination activity of RAD51 or its role in fork protection (**Figure 3A and B**). To distinguish the relative contributions of these RAD51 functions to MiDAS, we generated U2OS cells exogenously expressing RAD51 separation-of-function mutants, both of which have been shown to act in a dominant-negative manner. The RAD51 ATPase-dead K133R mutant forms highly stable RAD51 filaments which enhance RAD51-mediated protection of DNA but are deficient in HR (Mason et al., 2019; Morrison et al., 1999; Schlacher et al., 2011; Stark et al., 2002). Conversely, the Fanconi anaemia-associated RAD51 T131P mutant forms short RAD51 filaments which sustain HR but abrogate the protective function of RAD51 (Wang et al., 2015). U2OS cells constitutively expressing FLAG-tagged RAD51 variants, wild-type RAD51 (wt), RAD51 T131P or RAD51 K133R, exhibited similar cell cycle profiles following synchronisation (**Figure S3**), but the exogenous expression of the RAD51 mutants was noticeably lower compared to that of wt RAD51 (**Figure 3C**). This likely reflects the toxicity of the dominant-negative mutants when expressed at high levels under normal growth conditions. Interestingly, when exposed to mild replicative stress, cells expressing wt RAD51 or K133R, but not T131P, showed significant increases in mitotic EdU incorporation compared to parental U2OS cells (**Figure 3D-E**). We noticed, however, that cells expressing wt RAD51 also showed a significant increase in mitotic *γ*-H2AX foci, suggesting that increased wt RAD51 expression introduced an added source of genotoxic stress, leading to increased MiDAS (**Figure 3D and F**). Such an increase of mitotic *γ*-H2AX was not detectable in cells moderately expressing RAD51 K133R or T131P, suggesting that RAD51 K133R is uniquely able to facilitate mitotic EdU incorporation without affecting the level of under-replication to be dealt with. Collectively, these observations suggest that the enhanced DNA protection conferred by the RAD51 K133R mutant promotes MiDAS, even in the absence of proficient HR. In line with the proposed impact of RAD51 K133R on fork protection during interphase, RAD51 K133R specifically increased mitotic EdU foci which do not colocalise with *γ*-H2AX (**Figure 3G**).

**Figure 3:**
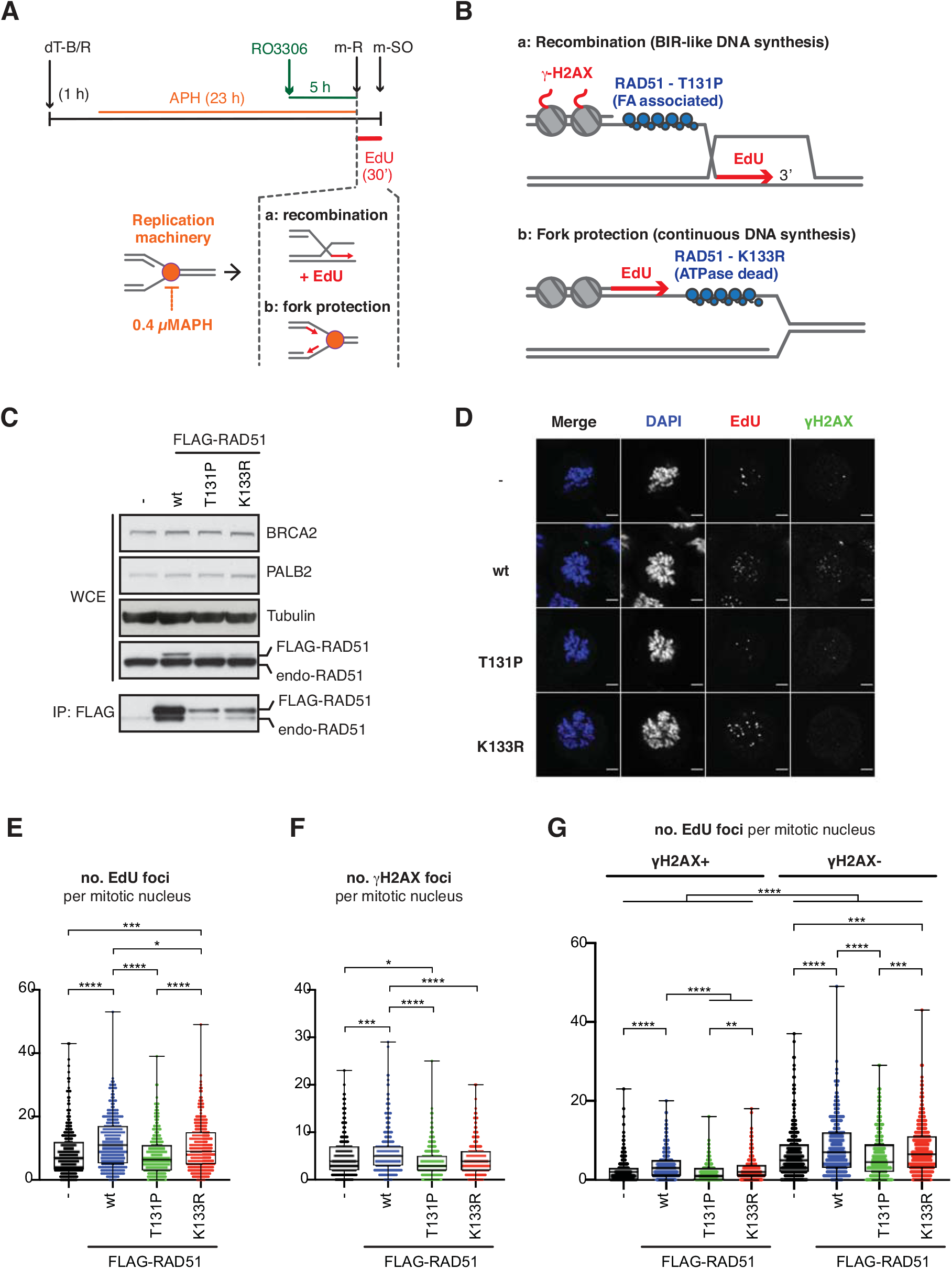
RAD51-mediated fork protection promotes non-break-associated MiDAS. **A.** Schematic diagram of experimental procedure, as in Figure 2A, except U2OS Flp-In T-REx cell lines constitutively expressing FLAG-tagged RAD51 separation of function mutants were used. Conceivable roles for RAD51 in MiDAS via break-associated recombination (a) or non-break-associated DNA synthesis (b) are also shown. **B.** Depiction of the expected RAD51 functions enabled by RAD51 T131P variant in catalysing recombination (a) and RAD51 K133R variant in protecting the replication fork (b). **C.** Western blot assessing BRCA2, PALB2, tubulin (loading control) and RAD51 protein levels in asynchronous U2OS Flp-In T-REx cells expressing the indicated FLAG-RAD51 variants. Anti-FLAG immunoprecipitate (IP) was probed for the presence of RAD51. **D.** Representative images of mitotic cells, stained for EdU and *γ*-H2AX, quantified in (E-G). Scale bar indicates 5 µm. **E-G.** Quantification of the number of EdU foci (E) and *γ*-H2AX foci (F) per mitotic nucleus. The data in (E) and (F) are quantified in (G) as the number of EdU foci that colocalise with *γ*-H2AX (*γ*-H2AX^+^ EdU foci) and the number of EdU foci that do not colocalise with *γ*-H2AX (*γ*-H2AX^-^ EdU foci) per mitotic nucleus. Data were obtained from three independent experiments (100 cells analysed per repeat); n=300 per condition. Data distribution is represented by box-and-whisker plots (whiskers spanning min-max range of values). Mann Whitney test p-values values are shown. Asterisks indicate p value ≤ 0.05 = *; ≤ 0.01 = **; ≤ 0.001 = ***; ≤ 0.0001 = ****.

### RAD51 S14 phosphorylation promotes MiDAS

Given that BRCA1 and BRCA2 functions in the DDR are downregulated in mitosis, RAD51 recruitment in mitosis is unlikely to occur via the canonical pathway mediated by the BRCA1-PALB2-BRCA2 complex. We hypothesised that RAD51 recruitment during mitosis instead relies on PLK1- and CK2-mediated RAD51 phosphorylation (**Figure 4A**). To test this, we exploited previously generated cell lines exogenously expressing untagged RAD51 phospho-mimetic (S14D) or phospho-null (S14A) mutants at similar levels, which were used to demonstrate the importance of RAD51 S14 phosphorylation for RAD51 recruitment to DSBs and stalled replication forks (Yata et al., 2014; Yata et al., 2012). Flow cytometric analyses of these cell lines revealed that exposure to mild replicative stress did not elicit pronounced changes in cell cycle progression for the S14A mutant compared to wt RAD51 expressing cells, while the RAD51 S14D mutant exhibited a faster progression into G2 phase (**Figure 4B and C**). These data suggest that the RAD51 S14A cell line experiences similar levels of under-replication compared to the wt RAD51 cell line, whilst the enhanced recruitment of RAD51 S14D mitigates the effects of APH on replication. When mitotic EdU incorporation was assessed (**Figure 4D**), the RAD51 S14A and S14D mutants exhibited significant reduction in mitotic EdU foci compared to wt RAD51, while no difference in the number of mitotic *γ*-H2AX foci was detected between the different RAD51 variant-expressing cell lines (**Figure 4E-G**). While reduced MiDAS in the RAD51 S14A cell line indicates a deficiency in MiDAS of the phospho-null mutant compared to wt RAD51, decreased MiDAS in the RAD51 S14D-expressing cell line likely reflects decreased under-replication experienced upon expression of the phospho-mimetic mutant, as indicated by its ability to progress through S-phase more rapidly in the presence of replication stress (**Figure 4C**). Collectively, these observations support the notion that RAD51 S14 phosphorylation promotes MiDAS in U2OS cells.

**Figure 4:**
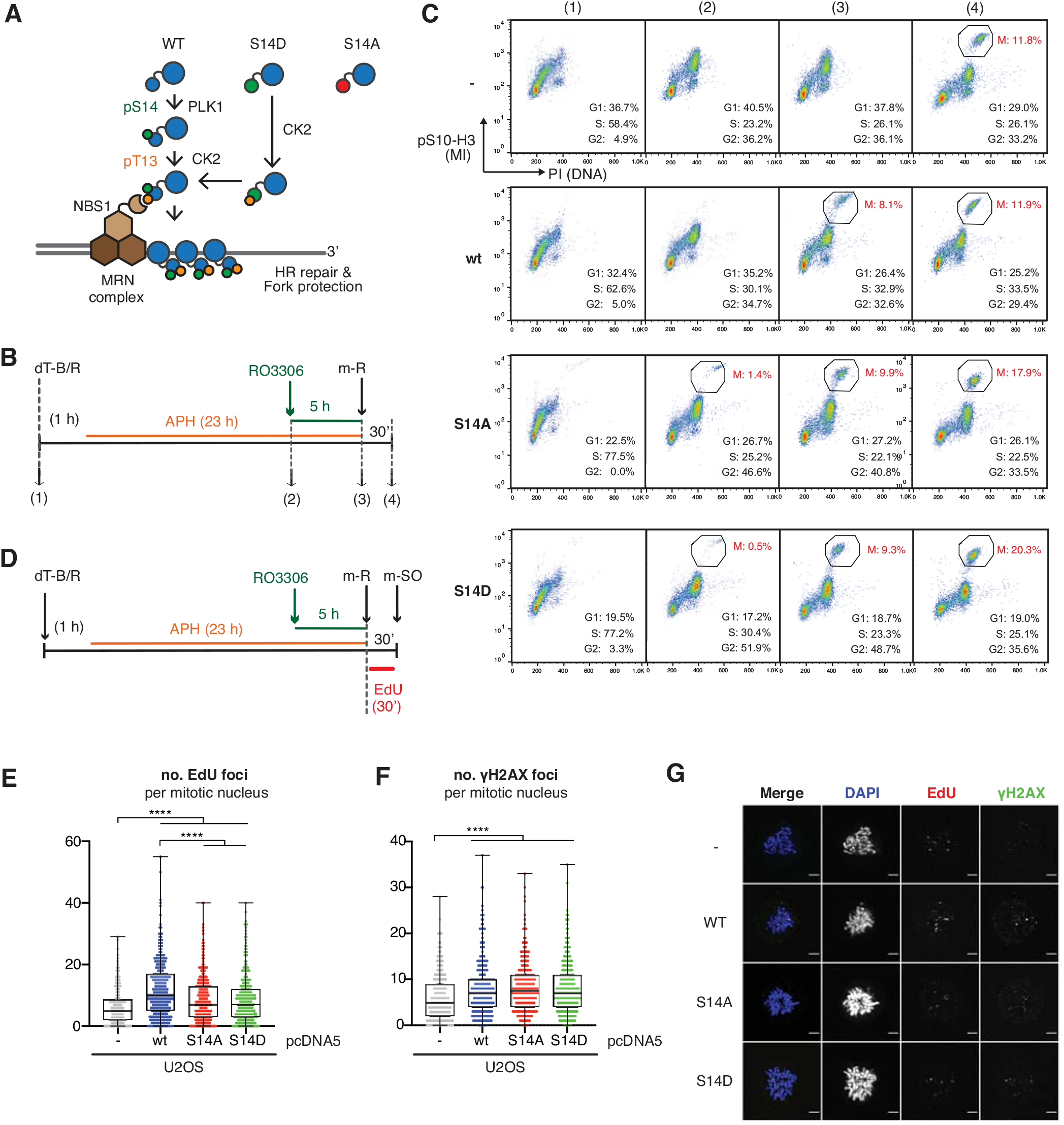
RAD51 S14 phosphorylation promotes MiDAS. **A.** Schematic illustration of the mechanism recruiting RAD51 to sites of DNA damage via PLK1-dependent RAD51 S14 phosphorylation, triggering CK2-dependent RAD51 phosphorylation at T13, which mediates NBS1 interaction. **B.** Schematic diagram of experimental procedure, as in Figure 3A, except U2OS cell lines stably expressing RAD51 S14 variants were used. **C.** FACS profiles of total cell populations at indicated time points shown in (B) for mitotic index (pS10-H3) and DNA content (PI). The G1, S and G2 populations in respective cell lines were estimated by the Watson Pragmatic algorithm (FlowJo), based on the PI staining of interphase cells. **D.** Schematic diagram of experimental procedure, as in (B), except cells were released into mitosis with 10 µM EdU exposure. **E.** Representative images of mitotic cells, stained for EdU and *γ*-H2AX, quantified in (F) and (G). Scale bar indicates 5 µm. **F-G.** Mitotic cells were collected by mitotic shake-off, stained and quantified for the number of EdU foci (F) and *γ*-H2AX foci (G) per mitotic nucleus. Data were obtained from three independent experiments (100 cells analysed per repeat); n=300 per condition. Data distribution is represented by box-and-whisker plots (whiskers spanning min-max range of values). Mann Whitney test p-values values are shown. Asterisks indicate p value ≤ 0.05 = *; ≤ 0.01 = **; ≤ 0.001 = ***; ≤ 0.0001 = ****.

### RAD51 phosphorylation is important for cellular proliferation

To better understand the importance of endogenous RAD51 phosphorylation for cellular proliferation, we set out to generate phospho-mimetic (S14D) or phospho-null (S14A or T13A-S14A) RAD51 mutants in U2OS cells by CRISPR/Cas9-mediated gene editing (**Figure S4A and B**). As PLK1- and CK2-phosphorylation of RAD51 is strictly sequential (no T13 phosphorylation is detected *in vivo* for the RAD51 S14A mutant) the T13A-S14A mutant and S14A mutant are functionally equivalent (Yata et al., 2012). Four homozygous RAD51 S14D mutants were obtained in the first round of screening as verified by sequencing (**Figure S4C and D**). No positive clones were obtained for the RAD51 S14A or T13A-S14A through two rounds of screening, suggesting the importance of S14 phosphorylation for U2OS cell viability (**Figure S4C**). Further attempts using other cell lines allowed us to obtain one heterozygous RAD51 S14A mutant in HEK293 cells, as verified by deep sequencing (**Figure S4E**). The absence of S14 phosphorylation in the CRISPR gene-edited phospho-mutants was confirmed by Western blot analysis (**Figure S4F**). The detection of major cell cycle markers revealed, however, that the HEK293 RAD51 S14A mutant exhibits increased expression of G1-S regulators, such as Cyclin E and p21, but reduced overall activity of PLK1, as detected by phosphorylation at T210 in the PLK1 activation T-loop. This observation suggested a disturbed cell cycle in the S14A mutant. We therefore complemented the HEK293 RAD51 S14A mutant by the stable expression of exogenous wt RAD51 (S14A+wt), which re-established RAD51 S14 phosphorylation (**Figure S4F**). Importantly, the above-mentioned cell cycle markers in the complemented RAD51 S14A+wt cell line remained unaffected, indicating that the RAD51 S14A mutant acquired secondary mutations during the cloning process, potentially to alleviate detrimental effects imposed by the loss of RAD51 T13/S14 phosphorylation.

Our evaluation of the CRISPR gene-edited cell lines confirmed that RAD51 S14D U2OS cells are more resistant than parental U2OS cells to DNA damage induction by the radio-mimetic neocarzinostatin (NCS) (**Figure S4G**). RAD51 S14D additionally displayed increased resistance to low dose of APH, as shown by clonogenic survival (**Figure S4H**) and cell division index (**Figure S4I**). These data are in line with the previously reported increase in RAD51 recruitment conferred by S14D mutation. In contrast, HEK293 RAD51 S14A, compared to the parental HEK293 cell line, displayed significantly increased sensitivity to low-dose APH, as shown by cell survival (**Figure S4J**) and cell division index (**Figure S4K**). Indeed, the RAD51 S14A mutation conferred a marked decrease in cell proliferation even in untreated conditions (**Figure S4K**). These phenotypes were partially rescued by wt RAD51 complementation in the RAD51 S14A+wt cell line (**Figure S4J-L**). HEK293 RAD51 S14A cells also exhibited decreased RAD51 foci formation and increased *γ*-H2AX compared to parental HEK293 cells in irradiated conditions (**Figure S4M and N**). RAD51 complementation in the S14A+wt cell line rescued IR-induced RAD51 foci formation, supporting the notion that the HEK293 RAD51 S14A mutant is defective in its recruitment to chromatin. The untreated HEK293 RAD51 S14A mutant also exhibited an increase in *γ*-H2AX foci compared to parental HEK293, suggesting that these cells experience high levels of endogenous replication stress, a phenotype that was rescued in the S14A+wt cell line. Taken together, these observations indicate that RAD51 S14 phosphorylation protects against replication stress and is important for the maintenance of proliferation in U2OS and HEK293 cells. It is important to note, however, that RAD51 complementation did not fully rescue the sensitivity to mild replication stress or reduced cell proliferation seen in the HEK293 S14A cell line. These observations suggest that this cell line is exposed to an additional source of endogenous genotoxic stress, different from the S14A mutation, potentially as a consequence of selective pressure due to the loss of S14 phosphorylation.

With these endogenous RAD51 mutant cell lines in hand, we then assessed them for mitotic EdU incorporation upon mild replicative stress. In line with the proficiency of RAD51 S14D mutant in protecting replication forks, we observed reduced mitotic *γ*-H2AX signal and mitotic EdU incorporation in the RAD51 S14D knock-in mutant compared to parental U2OS cells (**Figure S5A-E**). The RAD51 S14A knock-in mutant similarly exhibited reduced mitotic *γ*-H2AX signal as well as mitotic EdU incorporation compared to parental HEK293, and these phenotypes were partially rescued in the RAD51 S14A+wt cells (**Figure S5F-H).** We also observed impaired mitotic entry of the RAD51 S14A upon synchronisation with low-dose APH compared to parental HEK293 cells, although this phenotype was not rescued by exogenous wt RAD51 expression (**Figure S5I**). While we therefore cannot rule out the possibility that the observed mitotic phenotypes in these cell lines reflect a bias in the collection of severely under-replicated cells in mitosis, the rescue of mitotic EdU incorporation in RAD51 S14A+wt provided encouraging evidence for the importance of S14 phosphorylation for MiDAS in HEK293 cells.

### The RAD51 S14D mutant is resistant to spindle assembly checkpoint inhibition

MiDAS inhibition has been shown to be linked with an increase in chromosome mis-segregation and non-disjunction, indicating the detrimental consequences of mitotic division of an under-replicated genome (Minocherhomji et al., 2015). The timing of chromosome segregation is controlled by the spindle assembly checkpoint (SAC), which monitors the correct attachment of mitotic spindles to the kinetochore machinery assembled on the centromere of each sister chromatid. We hypothesised that SAC inhibition, by limiting the duration of early mitosis before anaphase onset, would aggravate the negative consequences of under-replication remaining in mitosis. To test this idea, cell survival of the endogenous RAD51 phospho-mutants was assessed upon exposure to an inhibitor of MPS1 kinase, a central component of SAC activation. A modest but significant increase in cell survival was observed for the U2OS RAD51 S14D cell line compared to the parental U2OS cell line upon exposure to the MPS1 inhibitor (**Figure 5A**). Clonogenic survival upon MPS1 inhibition demonstrated a clear increase in the RAD51 S14D mutant relative to parental U2OS cells (**Figure 5B**). Conversely, the HEK293 RAD51 S14A cell line exhibited reduced resistance to MPS1 inhibition compared to the parental HEK293 cell line, although this phenotype was not significantly rescued by wt RAD51 complementation (S14A+wt) (**Figure 5C**). Given that, compared to their respective parental cell lines, resistance against mild replicative stress was significantly increased in the U2OS RAD51 S14D cell line (**Figure S4H**) and reduced in both the HEK293 RAD51 S14A and S14A+wt cell lines (**Figure S4J**), these observations suggest that the sensitivity to SAC inhibition is correlated with cellular capacity to cope with replication stress.

**Figure 5:**
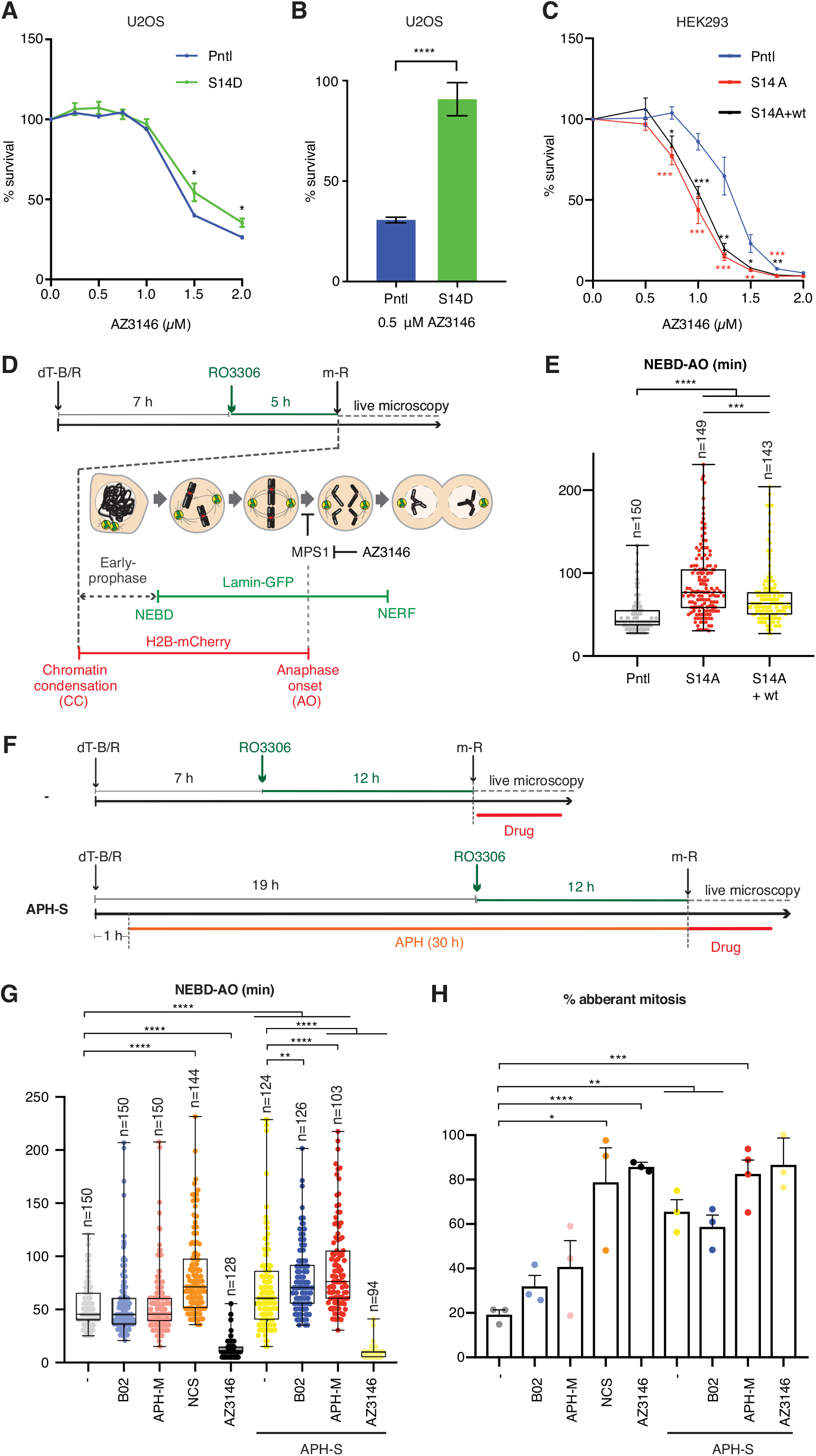
Inhibition of RAD51-mediated MiDAS delays anaphase onset in under-replicated cells. **A.** Cell survival of U2OS parental (Pntl) and RAD51 S14D knock-in mutant (S14D) upon exposure to AZ3146, as determined by WST-1 assay. Error bars represent SEM; n=12 from four independent experiments, including three technical repeats per experiment. Significant differences were assessed by unpaired t test. **B.** Clonogenic cell survival of U2OS parental (Pntl) and RAD51 S14D knock-in mutant (S14D) upon exposure to 0.5 µM AZ3146. Error bars represent SEM, n=7 from three independent experiments (at least two technical repeats were included per experiment). The significant difference between the samples was assessed by unpaired t test. **C.** Survival of HEK293 parental (Pntl), RAD51 S14A knock-in mutant (S14A) and wt RAD51 complemented (S14A+wt) cell lines upon exposure to MPS1 inhibitor (AZ3146), as determined by WST-1 assay. Error bars represent SEM; n=9 from three independent experiments, including three technical repeats per experiment. Significant differences compared to the parental HEK293 cell line were assessed by unpaired t test. **D.** Schematic diagram of experimental procedure for synchronising HEK293 cells exogenously expressing mCherry-tagged Histone H2B and GFP-tagged Lamin B1. Following dT-B/R and 5-hour RO3306 arrest, cells were released into mitosis (m-R) and mitotic progression was monitored by live microscopy. **E.** Mitotic duration of indicated HEK293 cell lines as measured from nuclear envelope breakdown (NEBD) to anaphase onset (AO). Pntl indicates the parental HEK293 cell line. Significant difference between samples were assessed by Mann Whitney test. **F.** Schematic diagram of experimental procedure, as in (D), except U2OS cells were optionally exposed to 0.4 µM APH (± APH-S), followed by 12-hour RO3306 arrest. Where indicated, mitotic progression was monitored in the presence of 20 µM RAD51 inhibitor (B02); 2 mM APH (APH-M), 2 µM MPS1 inhibitor (AZ3146), 50 nM neocarzinostatin (NCS) or vehicle (-). All drug treatments were added to cell culture directly after mitotic release (m-R). **G.** Mitotic duration of U2OS cells as measured from NEBD to AO. Cells that do not entry into anaphase within 4 hours from NEBD are excluded from analysis. Data was obtained from at least three independent experiments; n= the total number of cells monitored per condition. Data distribution is represented by box-and-whisker plots (whiskers spanning min-max range of values). Significant differences between samples were assessed by Mann Whitney test. **H.** Percentage of mitotic aberrations (lagging chromosomes, anaphase bridges) monitored during mitotic progression, shown in (H). Data represent the mean of at least three independent experiments, error bars represent SEM. Significant differences between samples were assessed by Mann Whitney test. For all statistical test, asterisks indicate p value ≤ 0.05 = *; ≤ 0.01 = **; ≤ 0.001 = ***; ≤ 0.0001 = ****.

### Anaphase onset is delayed under conditions of MiDAS inhibition

The observed correlation between cellular resistance against mild replication stress and SAC inhibition suggests the importance of SAC maintenance in allowing resolution of problems associated with under-replication prior to anaphase onset. This prompted us to consider whether the level of replicative stress and the proficiency of MiDAS could influence the progression of early mitosis. To test this idea, mitotic progression was monitored for synchronised cells stably expressing GFP-tagged Lamin B1 and mCherry-tagged Histone H2B, which permit the detection of nuclear envelope breakdown (NEBD) and anaphase onset (AO), respectively (**Figure 5D and S6A**). Live microscopy measurements of the time required from NEBD to AO revealed that HEK293 RAD51 S14A indeed significantly delayed anaphase onset compared to parental HEK293, which was partially but significantly rescued in the S14A+wt cell line (**Figure 5D and E, Supplemental Movies 1-3**). This trend was mirrored by the number of mitotic *γ*-H2AX foci, which was significantly increased in the RAD51 S14A cells, but reduced in the RAD51 S14A+wt cells (**Figure S6B and C**), while no significant difference in cell synchronisation was detected between the different HEK293 cell lines (**Figure S6D**). Given that HEK293 RAD51 S14A experiences increased endogenous replication stress and impaired MiDAS competency, which are partially rescued by exogenous wt RAD51 expression (**Figures S4J-N and S5F**), these data suggest that mitotic DNA damage resulting from endogenous replication stress and MiDAS defects contributes to the delay of anaphase onset.

We further examined whether mitotic duration of U2OS cells could be similarly affected by mild replicative stress. Indeed, low-dose APH in interphase (APH-S) delayed anaphase onset of U2OS cells (**Figure 5F and G, Supplemental Movies 4 and 5**), supporting the notion that replicative stress, arising from either endogenous or exogenous causes, impacts mitotic progression. This phenotype was further enhanced upon MiDAS inhibition by B02 or high-dose APH (APH-M) (**Figure 5G, Supplemental Movies 6-9**). Of note, U2OS cells exposed to low-dose APH exhibited increased aberrations of mitotic chromatin, such as lagging chromosomes and chromatin bridges, for which MiDAS impairment had little added impact, except that APH-M significantly increased mitotic cell death (**Figure 5H, S6E and S6F**). Regardless, anaphase onset was rapidly triggered by MPS1 inhibition in APH-S treated cells (**Figure 5G, Supplemental Movies 10 and 11**), suggesting that the SAC has a dominant role in mediating metaphase arrest in the presence of under-replication in mitosis. Importantly, we found that induction of mitotic DNA breaks upon NCS treatment similarly triggered a pronounced delay in anaphase onset (**Figure 5G, Supplemental Movie 1 2**). Altogether, given that MiDAS inhibition by mitotic B02 or APH-M treatment induces mitotic DNA damage (**Figure 2G**), these observations support the notion that the generation of mitotic DNA breaks and/or the persistence of under-replicated DNA prevent SAC satisfaction.

### Replication stress affects centromeric integrity in mitosis

The SAC is satisfied when the centromeres of sister chromatids attach to opposite poles of the spindle, such that under-replicated centromeres potentially delay anaphase onset simply due to the failure of this process. Additionally, it has been demonstrated that mitotic DNA damage at centromeres, but not at chromosome arms, prevents SAC satisfaction (Mikhailov et al., 2002). Centromeres have also been shown to exhibit high levels of sister-chromatid exchange (cen-SCE), as detected by chromosome-orientation FISH (CO-FISH), supporting the idea that centromeres are prone to breakage and/or form aberrant DNA structures under certain physiological conditions such as in cancer cells and during replicative senescence (Jaco et al., 2008; Giunta and Funabiki, 2017) (**Figure 6A**). We therefore wondered whether centromeric DNA structure might be affected by mild replicative stress and MiDAS inhibition in a manner that prevents satisfaction of the SAC. Indeed, our CO-FISH analyses of U2OS cells exposed to mild replication stress (APH-S) showed an increase in aberrant centromeric signal, although this phenotype was not rescued upon MiDAS inhibition by mitotic APH (APH-M) treatment (**Figure 6B-D**). This observation suggests that, while centromeres undergo cen-SCE upon exposure to mild replication stress, this occurs in interphase and does not entail MiDAS (**Figure 6D**). Unexpectedly, we found a significant increase in aberrant centromeric signal in cells exposed to low-dose APH followed by mitotic RAD51 inhibition (B02) (**Figure 6D**), potentially reflecting enhanced distortion of centromeric DNA structure in the absence of RAD51-mediated protection in mitosis. Interestingly, while analysing CO-FISH signals, we noticed that their intensities were also increased significantly in cells exposed to APH-S, a phenotype that was further exacerbated by mitotic B02- or APH-M treatments (**Figure 6E**). Given that FISH signal intensity reflects the presence of ssDNA to which the FISH probes hybridise, these observations suggest that MiDAS inhibition increased centromere ‘fragility’, such that an increased level of ssDNA within the centromeric DNA was exposed, either in the context of native centromeric DNA or during the process of sample preparation. Overall, these data suggest that mild replication stress induces distortion of centromeric DNA structure in mitosis, the formation of which is alleviated by MiDAS.

**Figure 6:**
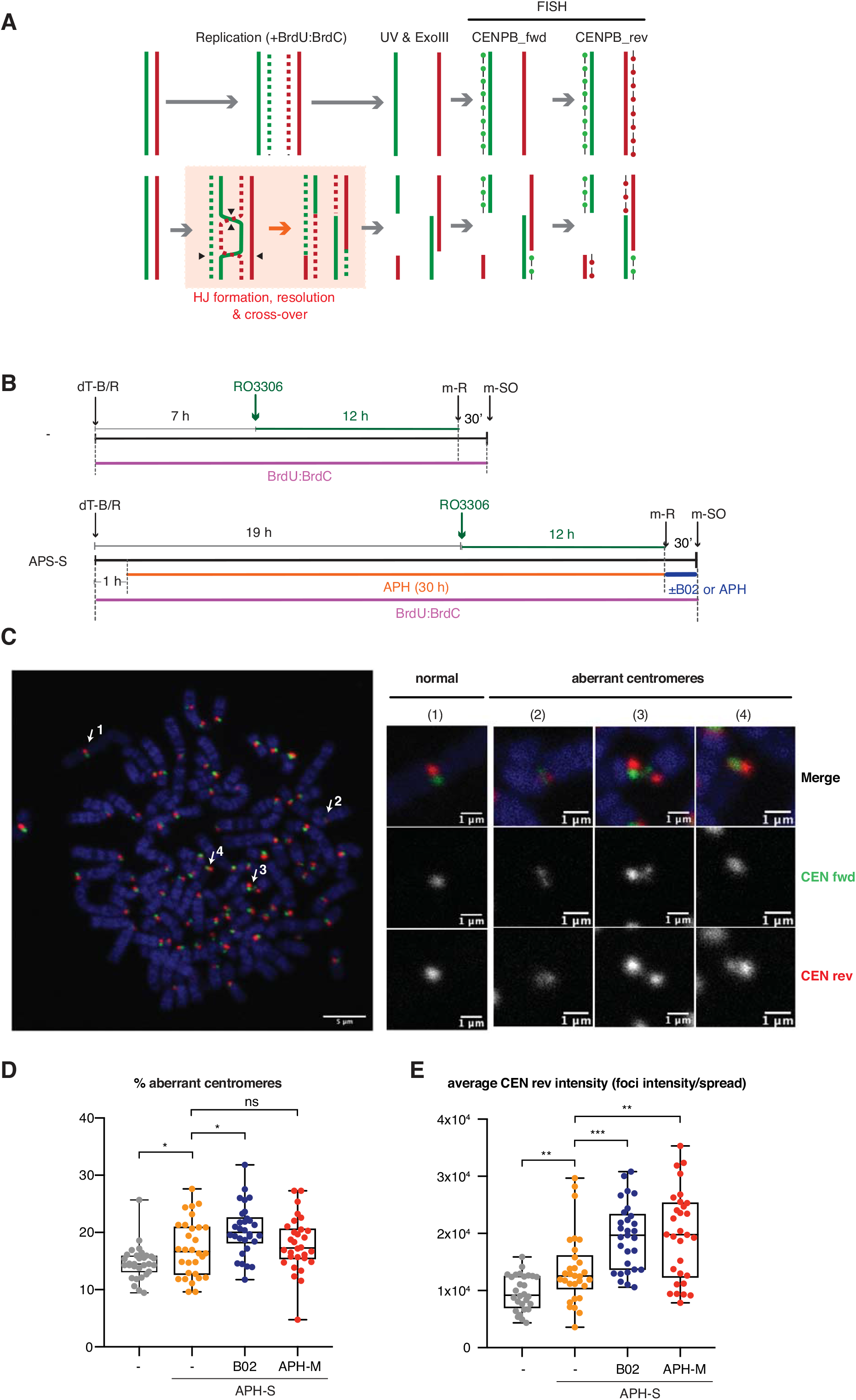
RAD51 inhibition in mitosis de-protect centromeric DNA. **A.** Schematic diagram for the experimental procedure of CO-FISH. Newly synthesised DNA incorporates BrdU, which is then digested by UV and ExoII treatment. Forward (fwd) and reverse (rev) FISH probes, which hybridise to centromere-specific CENPB-box sequences of the undigested strand, were hybridised in this order. **B.** Schematic diagram showing U2OS cell synchronisation, as in Figure 5F, except cells were exposed to BrdU as cells were released from thymidine block to allow digestion of newly synthesised DNA during the CO-FISH procedure. Cells were then released into mitosis with or without 20 µM RAD51 inhibitor (B02); 2 µM aphidicolin (APH (M)) or vehicle (-). **C.** Representative image of CO-FISH. Arrows indicate examples of normal centromere signal (1) and aberrant centromere signal (2-4). **D.** Percentage of aberrant centromere signal per metaphase spread obtained for U2OS cells. **E.** Same data as (D), with the average focus intensity of the reverse centromere probe (CEN-rev) quantified per metaphase spread. (D-E) Data were obtained from three independent experiments (10 cells analysed per repeat); n=30 per condition. Data distribution is represented by box-and-whisker plots (whiskers spanning min-max range of values). Mann Whitney test p-values are shown. In panel (D) unpaired t test p-values values are shown. Asterisks indicate p value ≤ 0.05 = *; ≤ 0.01 = **; ≤ 0.001 = ***; ≤ 0.0001 = ****.

## Discussion

### Identification of RAD51 as a new player in MiDAS

This study demonstrates a previously unrecognised mitotic role for RAD51 in resolving under-replicated mitotic chromatin. The role of RAD51 in MiDAS was assessed upon siRNA-mediated RAD51 depletion; addition of a small molecule inhibitor (B02); and in cells expressing RAD51 separation of function mutants (T131P and K133R) or phospho-mutants (S14A and S14D) (**Figures 1**-**4, S5**). In general, the importance of RAD51 in interphase complicates the analysis of its role in MiDAS, as this influences the extent of under-replication that passes into mitosis. In this context, the acute inhibition of RAD51 by B02 offered the most reliable method by which the role of RAD51 in mitosis could be studied without affecting the previous interphase. B02 treatment of mitotic cells drastically reduced mitotic EdU incorporation while increasing mitotic *γ*-H2AX and RPA foci (**Figure 2F-H**), indicating a direct role for RAD51 in promoting MiDAS, involving the protection of stalled replication forks from DNA resection and collapse. Given that B02 specifically inhibits *de novo* recruitment of RAD51, our findings indicate that RAD51 is actively recruited in mitosis. This is in line with the previous observation that RAD51 is removed by RECQ5 from under-replicated CFSs prior to mitotic entry (Di Marco et al., 2017). Our study also suggests that *de novo* RAD51 recruitment plays a role at centromeres, which are enriched for PLK1 (Arnaud et al., 1998). We propose that RAD51 is recruited on mitotic chromatin in a PLK1-dependent manner to protect stalled replication forks from mitotic nucleolytic attack, potentially mediated by MUS81 and MRE11, hence allowing for the completion of DNA replication and subsequent cell division (**Figure 7**).

**Figure 7:**
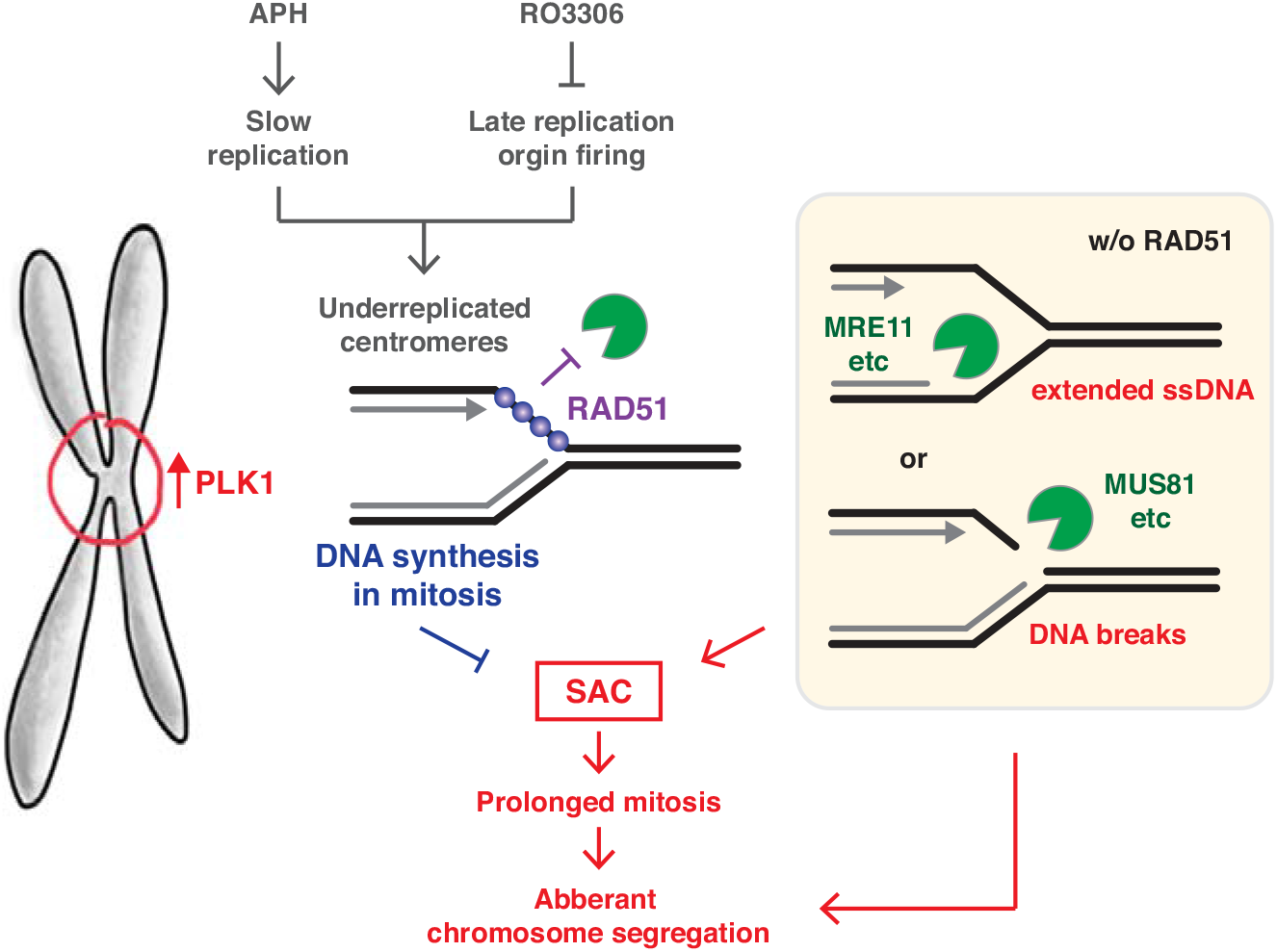
Model. Mild replicative stress (i.e., 0.4 µM aphidicolin) and the suppression of late origin firing (i.e., RO3306) may result in substantial under-replication, which is undetected by the G2/M checkpoint. As cells move into mitosis, RAD51 is recruited to replication forks in a manner dependent on PLK1, and protects them from nucleolytic attack, potentially mediated by MUS81 and MRE11, allowing the completion of DNA replication. Centromeric under-replication and subsequent MiDAS or mitotic DNA damage prevents SAC inactivation. RAD51-mediated protection of centromeric DNA structure, which are enriched with PLK1, ensures appropriate chromosome segregation.

Previous studies have shown that the DDR is largely attenuated in mitosis, such that the recruitment of various DNA repair proteins is inhibited during this phase of the cell cycle. It is important to note, however, that the early DDR remains intact and the MRN complex is efficiently recruited to mitotic DNA breaks (Giunta et al., 2010; Orthwein et al., 2014). This supports the idea that RAD51 recruitment can be achieved in mitosis by PLK1-dependent RAD51 S14 phosphorylation, which leads to its direct interaction with the MRN complex (Yata et al., 2012). This notion is in agreement with a previous study showing an active role of PLK1 in promoting MiDAS (Minocherhomji et al., 2015). PLK1-mediated RAD51 S14 phosphorylation is also promoted by TOPBP1, which has likewise been shown to promote efficient DNA synthesis in mitosis (Moudry et al., 2016; Pedersen et al., 2015). Therefore, it is conceivable that both PLK1 and TOPBP1 impact MiDAS, at least in part, through the recruitment of RAD51. It is noteworthy that loss of endogenous RAD51 S14 phosphorylation in CRISPR gene-edited phospho-mutants greatly affected cell cycle progression and cellular proliferation (**Figures S4K and S5I**), indicating that this is a major pathway for RAD51 recruitment even under unperturbed conditions. It is likely that the HEK293 RAD51 S14A knock-in acquired secondary mutations as an adaption to the loss of RAD51 phosphorylation. For instance, cyclin E overexpression in HEK293 RAD51 S14A cells might drive increased origin firing as a response to the replication stress introduced by the loss of PLK1-dependent RAD51 recruitment (**Figure S4F**). While, similarly to RAD51 depletion, the impact of the S14 knock-in mutations on cell cycle progression ultimately impedes the comparative analyses of MiDAS across these cell lines, it is interesting that an RAD51 S14A mutant could not be obtained in U2OS cells, in which the majority of MiDAS is not associated with *γ*-H2AX, so is likely break-independent. It is tempting to speculate that the phospho-null RAD51 mutation is better tolerated in HEK293 cells, in which break-induced MiDAS was more common, such that RAD52 function may compensate for the lack of RAD51 recruitment (see below). The importance of RAD51 phosphorylation in different cell lines warrants further investigation.

### Mechanism mediating MiDAS

Previous studies of MiDAS have focused on CFSs and telomeres, where MiDAS occurs via a RAD52-dependent break-induced mechanism (Bhowmick et al., 2016; Min et al., 2017). However, our study illuminated the diverse nature of MiDAS depending on the cell lines and specific genomic loci analysed, as well as the experimental conditions used during cell synchronisation. In particular, our analyses revealed that the majority of MiDAS events in U2OS cells occur independently of DNA breaks, as indicated by the lack of *γ*-H2AX at sites of mitotic EdU incorporation (**Figures S1G-H and 2I**). Interestingly, while BIR is proposed to occur via conservative replication, a recent study reports that the majority of MiDAS events in U2OS are semi-conservative, supporting the view that MiDAS in U2OS is not entirely a result of BIR (Chappidi et al., 2020). In contrast, mitotic EdU foci in HEK293 cells largely colocalise with *γ*-H2AX (**Figure S1H**). Indeed, compared to U2OS, HEK293 cells exhibited a high basal level of *γ*-H2AX, suggesting that break-induced MiDAS is more prevalent in cells that experience greater replication stress (**Figure S4F**). We envisage that break-independent MiDAS occurs at stalled forks which retain the DNA replication machinery and can therefore continue DNA synthesis into mitosis. While the extension of G2 phase by CDK1 inhibition significantly decreases mitotic EdU incorporation (Garribba et al., 2018), a substantial amount of MiDAS was detectable even after a prolonged CDK1 inhibition of 12 hours (**Figure S1E**). This observation supports the notion that the replication of certain difficult-to-replicate loci requires CDK1 activity or indeed, mitotic entry.

Several events occur concomitantly with mitotic entry which may promote the completion of DNA replication. These proposed mechanisms include CDK1-mediated activation of nucleases such as MUS81 to trigger break-induced MiDAS and WAPL-mediated removal of cohesin (Minocherhomji et al., 2015). Interestingly, WAPL is also proposed to promote RAD51-dependent replication restart at broken forks (Benedict et al., 2020). We also envisage that the removal of several chromatin-associated proteins in interphase, such as the transcriptional machinery and the cohesin complex at chromosome arms, may prepare the chromatin environment to accommodate MiDAS. Additionally, given that CDKs trigger replication origin firing by phosphorylating the CMG (Cdc45-MCM-GINS) helicase complex, it is also plausible that fully activated CDK1 in mitosis may assist late or dormant origin firing in under-replicated regions which retain the CMG complex. In support of this notion, enriched chromatin association of the CMG complex components in cells exposed to mild replicative stress has been observed (Minocherhomji et al., 2015). Notably, we observed that WEE1 inhibition, which activates CDK1, increased mitotic EdU incorporation (**Figure 1E**). While this may simply reflect the mitotic entry of severely under-replicated cells, no significant increase in mitotic *γ*-H2AX or RPA foci was detectable upon WEE1 inhibition (**Figure 1F-G**), perhaps indicating that enhanced CDK1 activity *per se* promotes break-independent MiDAS.

While our study agrees with the previous observation that RAD52 promotes MiDAS via a break-induced mechanism, we also found that this constitutes a relatively small subset of MiDAS events in U2OS cells (**Figure 2F, I and S2F, H**). In contrast, RAD51 appears to be essential for a broader range of MiDAS events, which take place with and without DNA breakage. Considering that the DNA structure-specific nuclease activities are at their highest during mitosis, the protection of DNA intermediates arising from MiDAS is likely vital. Indeed, MUS81 can cleave recombination intermediates formed during BIR, as well as ongoing replication forks, as shown in interphase (Duda et al., 2016; Matos et al., 2011; Mayle et al., 2015; Wyatt et al., 2013). Therefore, RAD51-mediated protection against nucleolytic cleavage may be a common requirement for both break-induced and break-independent MiDAS. Notably, bright RAD51 foci, which are typically associated with damage-induced HR repair, were not detectable in early mitosis. This lack of RAD51 foci is consistent with an early mitotic role for RAD51 in replication fork protection, as the protective role of RAD51 at stalled forks is not associated with formation of foci (Petermann et al., 2010). This notion is further supported by our observation that expression of exogenous RAD51 K133R, which confers enhanced fork protection but is HR-defective, increased mitotic EdU incorporation that did not colocalise with *γ*-H2AX (**Figure 3G**). It is, nonetheless, noteworthy that, while *γ*-H2AX-associated MiDAS was not significantly affected by the K133R mutation, B02-mediated mitotic RAD51 inhibition clearly reduced it (**Figure 2I**). Indeed, during the preparation of this manuscript, RAD51 was shown to catalyse break-associated MiDAS at the Fragile X locus, a folate-sensitive rare fragile site (Garribba et al., 2020). Therefore, while BIR-like MiDAS is also RAD51-dependent, it is surprising that the K133R mutation, which affects both the HR and protective functions of RAD51, does not appear to affect break-associated MiDAS. This might be best explained by a dual, opposing effect of the K133R mutation, where the lack of HR reduces break-induced MiDAS, but the enhanced protection of branched DNA structures promotes it. Of note, RAD51 T131P expression did not significantly affect mitotic EdU incorporation compared to that seen in the parental cell line (**Figure 3E**). This suggests that the expected loss of RAD51-mediated DNA protection is not exhibited at such low levels of T131P expression. Indeed, T131P cells surprisingly exhibited decreased mitotic *γ*-H2AX foci, further refuting the loss of DNA protection in this cell line (**Figure 3F**).

### The spindle assembly checkpoint (SAC) senses under-replication in mitosis

Our study also highlighted the importance of the SAC, which appears to be related to the amount of replication stress experienced by the cell. One intriguing possibility is that the SAC may ‘sense’ mitotic DNA damage or under-replication, thereby preventing SAC silencing until MiDAS has been completed. In line with this idea, we found that APH-induced replication stress during interphase significantly delayed anaphase onset in U2OS cells, and that this delay was dependent on the SAC (**Figure 5G**). The APH-induced delay in mitotic progression was further increased upon MiDAS inhibition by either mitotic RAD51 inhibition or mitotic APH treatment. Altogether, these findings suggest that the inability to complete replication in mitosis and/or the persistence of mitotic DNA breaks prevents SAC inactivation.

The notion that under-replicated DNA or mitotic DNA breaks affect SAC signaling led us to further consider whether these mitotic DNA stresses occur at the centromere. Curiously, mitotic EdU incorporation or *γ*-H2AX were rarely detectable at centromeres under the conditions we employed, potentially suggesting that centromeric DNA synthesis is extremely inefficient and/or the DDR is downregulated in mitosis at these regions. However, we noticed that the conditions of MiDAS inhibition led to an increase in centromeric FISH signal intensity, which we propose is an early indication of centromeric ‘fragility’ (**Figure 6E**). Mitotic RAD51 inhibition further triggered an increase in aberrant centromeric CO-FISH signal in cells that experienced mild replication stress (**Figure 6D**). Considering the role of RAD51 in protecting under-replicated mitotic chromatin and the fact that centromeres, being highly repetitive and transcriptionally active even during mitosis, are among the most difficult-to-replicate regions, we hypothesise that centromeres are deprotected in the absence of RAD51 and thereby become particularly prone to form non-B DNA structures. The increased centromeric instability observed upon MiDAS inhibition presumably affects kinetochore integrity and microtubule binding, thereby preventing satisfaction of the SAC. It is tempting to speculate that the inherent vulnerability of the centromere to under-replication enables it to act as a gauge for overall genome replication and hence maintains SAC activation to maximise genome replication prior to anaphase. This might be particularly advantageous for organisms with large genomes and may, in part, explain the complexity of the centromere in eukaryotes.

### Implications for cancer therapy

The identification of players in the MiDAS pathway may reveal promising targets for cancer therapy. This study reveals a novel role for RAD51 in promoting both break-induced and break-independent MiDAS mechanisms. Furthermore, RAD51 phosphorylation appears to mediate RAD51 recruitment in mitosis. A better understanding into how PLK1-dependent RAD51 phosphorylation is regulated upon DNA damage or replication stress may reveal druggable targets to abrogate RAD51 recruitment in mitosis, and thereby inhibit MiDAS. Lastly, inhibition of SAC activity was shown to severely decrease cell survival in cells that exhibit high levels of replication stress. This suggests that therapies aimed at increasing cellular replication stress may be particularly effective when combined with chemotherapeutic treatments targeting the spindle checkpoint.

## Acknowledgements

We thank Profs Timothy C. Humphrey, Ian D. Hickson, Chris J. Norbury, Dr James Carrington, and members of the Esashi laboratory for helpful discussions; Drs Nigel Rust and Michal Maj for assistance with FACS; and Dr Alan Wainman for assistance with microscopy. FE was supported by the Wellcome Trust Senior Research Fellowships in Basic Biomedical Science (101009/Z/13/Z), and is thankful for supports from the Edward Penley Abraham Research Fund. IEW was a recipient of the Medical Sciences Graduate School studentship, funded by the Medical Research Council (14/15_MSD_439771). XS receives the Oxford Cancer Centre Cancer Research UK D.Phil studentship (CRUK-OC-DPhil17-XS). LR is a recipient of the Oxford Interdisciplinary Bioscience Doctoral Training Partnership, sponsored by the Biotechnology and Biosciences Research Council (BB/M011224/1, Project 1757783). AB is thankful for the support from the departments of Pathology, Biochemistry, Pharmacology and Physiology, Anatomy and Genetics and from the John Fell Fund and Wellcome ISSF.

## Author contributions

F.E. and I.E.W. conceived and planned the project; I.E.W. conducted all experiments unless otherwise specified; F.E., I.E.W. and L.R. generated CRISPR knock-in mutants with the help of C.R. and A.B; F.E. performed most Western blots, except Fig 1B and Fig S2E, which were conducted by X.S. and I.E.W., respectively; L.R. contributed to the identification of cell division index; X.S. contributed to the installation of the cen-CO-FISH methodology; F.E. and I.E.W. wrote the manuscript with input from all contributing authors.

## Declaration of Interests

We declare no financial and non-financial conflict of interests on this study.

## Methods

### Cell culture

All cell lines were cultured at 37 °C with CO_2_ in Dulbecco’s modified Eagle medium supplemented with 10% v/v fetal bovine serum, streptomycin (0.1 mg/ml) and penicillin (100 units/ml). U2OS Flp-In T-RE-x cell lines expressing RAD51 variants (**Figures 3 and S4**) were generated by transfecting pcDNA5/FRT encoding FLAG-tagged RAD51 variant to U2OS Flp-In T-REx (a kind gift from Daniel Durocher), followed by hygromycin selection at 200 µg/ml. All transfections were performed using JetPrime (Polyplus transfection). All cell lines used to assess mitotic progression (**Figures 5 and S7**) were generated by co-transfecting pcDNA3 encoding mCherry-tagged Histone H2B (Addgene: 20972) and pcDNA3 encoding GFP-tagged Lamin B1 (a kind gift from David Vaux). Stable clones were isolated by FACS. U2OS RAD51 variant expressing cells (**Figure 4**) were previously generated (Yata et al., 2012). To generate RAD51 S14 knock-in mutants (**Figures S5 and S6**) cells were co-transfected with a CRISPR/Cas9 construct (pSpCas9(BB)-2A-GFP) expressing guide RNA targeting the RAD51 S14 locus (AGCAAATGCAGATACTTCAGTGG), as well as the relevant ssDNA repair template (**Table S1**). Clones were confirmed by DNA sequencing of the RAD51 S14 locus and immunoblotting for RAD51 S14 phosphorylation. The wild-type complemented HEK293 RAD51 S14A mutant was generated by co-transfecting pcDNA5/FRT encoding wild-type RAD51 and pcDNA-DEST26 (Invitrogen), followed by single cell sorting (FACS) and G418 selection at 400 µg/ml. MUS81 was depleted using siMUS81 (ID: 130843, ThermoFisher Scientific); RAD52 was depleted using ONTargetplus Human RAD52 siRNA (Horizon) and RAD51 was depleted using a mixture of two siRNAs as previously published (Yata et al., 2012). MISSION siRNA Universal Negative Control #1 (Sigma-Aldrich) was used as a negative control. All siRNA treatments were performed with 20 nM siRNA using JetPrime (Polyplus transfection).

### Chemical inhibitors

Where indicated, cells were treated with aphidicolin (Santa Cruz); RO3306 (Cayman Chemical); B02 (Cayman Chemical); AICAR (Sigma-Aldrich); AZD1775 (Selleckchem); AZ3146 (Santa Cruz); Neocarzinostatin (Sigma-Aldrich) at the indicated concentrations and for the indicated lengths of time.

### Cell survival assay

For clonogenic survival assay, U2OS cells were plated at a density of 40 or 100 cells per well in a 6-well plate. Once adhered, the corresponding drug was added to the cell culture at the indicated dose and cells were incubated for 8 days. Cells were then fixed and stained in 50% methanol, 7% acetic acid and 0.1% Coomassie Brilliant blue. Colonies of > 50 cells were counted, and percentage survival was calculated relative to the plating efficiency of the vehicle-treated control. For WST-1 cell proliferation assay, cells were seeded at a density of 5000 cells (U2OS) or 10 000 cells (HEK293) per well in a 96-well plate. Once adhered, the corresponding drug was added to the cell culture at the indicated dose and cells were incubate for 5 days. Cell survival relative to vehicle-treated cells was assessed using the WST-1 kit (Roche) according to the manufacturer’s instructions.

### Extract preparation, Western Blotting, Immunoprecipitation

Cells were collected by trypsinization, washed with ice-cold in PBS and incubated on ice in extraction buffer (50 mM Tris-HCl pH 8.0, 150 mM NaCl, 2 mM EDTA pH 8, 0.5% NP40, 10 mM Benzamide Hydrochloride, 20 mM NaF, 20 mM *β*-glycerophosphate, 1 mM Na_3_VO_4_, 5 mM MgCl_2_, 125 U/ml Benzonase nuclease (Novagen) and 1x Protease inhibitor cocktail (Sigma-Aldrich) for at least 30 min. Supernatant was collected as whole cell extract. For immunoprecipitation, the obtained cell extracts were pre-cleared with uncrosslinked Affiprep Protein-A beads (BioRad) for at least 1 hour at 4 °C. The pre-cleared lysate was subsequently incubated with antibody-crosslinked Affiprep Protein-A beads overnight at 4 °C. After extensive washing in extraction buffer, immune complexes were eluted from the beads in NuPAGE LDS sample buffer (ThermoFisher Scientific) supplemented with 12 mM DTT by heating the sample at 85 °C for 5 min. Western blotting was performed following standard protocol. Primary and secondary antibodies were applied at the dilutions described in the Supplemental Methods. Where indicated, the membrane was treated with Re-Blot Plus Mild Solution (Millipore) before incubating with another antibody.

### Flow cytometry

Cells were collected by trypsinization and fixed in 70% ethanol at approximately 10^6^ cells/ml. After fixation, cells were permeabilised in PBS with 0.1% Triton X-100 (Sigma-Aldrich) and 1% BSA (Sigma-Aldrich) for 15 min. To detect mitotic cells, the permeabilized cells were incubated for 1 hour with anti-phospho-S10 Histone H3 (06-570, Merck Millipore) in PBS containing 0.1% Tween-20 (Sigma-Aldrich) and 1% BSA, followed by a subsequent incubation with Alexa Fluor 488-conjugated secondary antibody (ThermoFisher Scientific) for 30 min. Cells were counter-stained in PBS with 0.1% BSA, 0.1 mg/ml RNAse A (Sigma-Aldrich) and 2 µg/ml propidium iodide (Sigma-Aldrich) for 30 min. Cell cycle distribution was analysed using a FACSCalibur (Becton Dickinson) and FlowJo analysis software (version 10.6.2). Cell counts in G1, S, G2 were estimated using the Watson Pragmatic algorithm in FlowJo on gated interphase cells and combined with quantification of gated mitotic cells to calculate the indicated cell cycle distributions.

### Cell synchronisation

For double thymidine block-and-release, cells were cultured in the presence of 2 mM thymidine (Sigma-Aldrich) for 18 hours and released in fresh medium for 9 hours, before being incubated in 2 mM thymidine for an additional 17-18 hours. Cells were released into fresh medium to allow for progression through S-phase. Where indicated, 0.4 µM aphidicolin was added 1 hour upon release from the thymidine block. Cells were subsequently arrested in G2 upon the addition of 9 µM RO3306 at the indicated time points and incubated for 5 or 12 hours. Cells were released into fresh medium to allow for mitotic entry and collected by mitotic shake-off after 30 min. EdU pulse-labelling was performed at the indicated time points by 30 min incubation in the presence of 10 µM 5-ethynyl-2-deoxyuridine (Life Technologies).

### Immunofluorescence (IF)

When detecting IF upon irradiation-induced DNA damage, cells were seeded on coverslips one day prior. Where indicated, a Gravitron RX 30/55 (Gravatom) was used to irradiate cell cultures at the stated dose. Cells were returned to cell culture for 3 hours post-irradiation and subsequently fixed and permeabilised for 20 min in PTEMF buffer (20 mM PIPES pH 6.8, 10 mM EGTA, 0.2% Triton X-100, 1 mM MgCl_2_, 4% paraformaldehyde). Primary and secondary antibodies were applied at the dilutions listed in table S2 and incubated at room temperature for at least 1 hour. When detecting IF in mitotic shake-off, mitotic cells were seeded onto poly-L-lysine coated coverslips and allowed to adhere for 5-10 min before being fixed and permeabilised for 20 min in PTEMF buffer. Prior to antibody incubation, fixed and permeabilized cells were incubated for 1 hour in PBS containing 2 mM CuSO4, 10 mM sodium ascorbate and 10 µM Alexa Fluor azide (ThermoFisher Scientific) to fluorescently label incorporated EdU. Coverslips were washed in PBS with 0.5% Triton X-100 in between incubations and mounted in DAPI-containing ProLong Gold antifade reagent (Life Technologies).

### Metaphase spreads and CO-FISH

Cells were synchronised by double thymidine block and released into S phase in the presence of 7.5 µM BrdU and 2.5 µM BrdC. Where indicated, 0.4 µM aphidicolin was added 1 hour upon release from the thymidine block. Cells were subsequently arrested in G2 upon the addition of 9 µM RO3306 at the indicated time points and incubated for 12 hours. Cells were released into fresh medium containing 0.1 µg/ml colcemid (Millipore), 7.5 µM BrdU and 2.5 µM BrdC for 30 min. Mitotic cells were subsequently collected by mitotic shake-off and swollen in 0.56% w/v KCl for 20 min at 37 °C. Cells were fixed in methanol:acetic acid (3:1) and dropped onto glass slides. The metaphase spreads were prepared for CO-FISH according to the previously published protocol (Giunta and Funabiki, 2017), as outlined in the Supplemental Methods.

### Microscopy analysis and live cell imaging

For fixed samples, all images were captured using an Olympus FV1000 Laser Scanning Microscope with Becker and Hickel FLIM system and Olympus imaging software. Foci counting was performed using FIJI (Image J version 2.0.0-rc-65/1.52a) and the GDSC (FindFoci) ImageJ plugin. Live cell images were captured using a Zeiss 880 inverted confocal microscope. Images were captured every 4.5 min (**Figure 6E**) or 5 min (**Figure 6G**) for at least 6 hours.

### Statistical analysis

All statistical analysis was performed using GraphPad PRISM 7 For Mac OS X.

## Notes

### Competing Interest Statement

The authors have declared no competing interest.

