## Supplemental Information for "The RAD51 recombinase protects mitotic chromatin in human cells"

### Index

#### Supplemental Figures

**Figure S1:** Evaluation of DNA synthesis upon mild replicative stress in mitosis.

**Figure S2:** Evaluation of RAD51 inhibitor B02 and a comparative analysis between RAD51 and RAD52 depletion for MiDAS.

**Figure S3:** Characterisation of U2OS cells expressing RAD51 separation-of-function mutants.

**Figure S4:** Evaluation of endogenous RAD51 S14 missense mutant cells.

**Figure S5:** Analysis of MiDAS in CRISPR gene-edited RAD51 S14 knock-in mutants.

**Figure S6:** Impact on RAD51 defects in mitotic chromosome integrity.

#### Supplemental Figure Legends

#### Supplemental Movie Legends

#### Supplemental Methods

Wassing et al., Figure S1

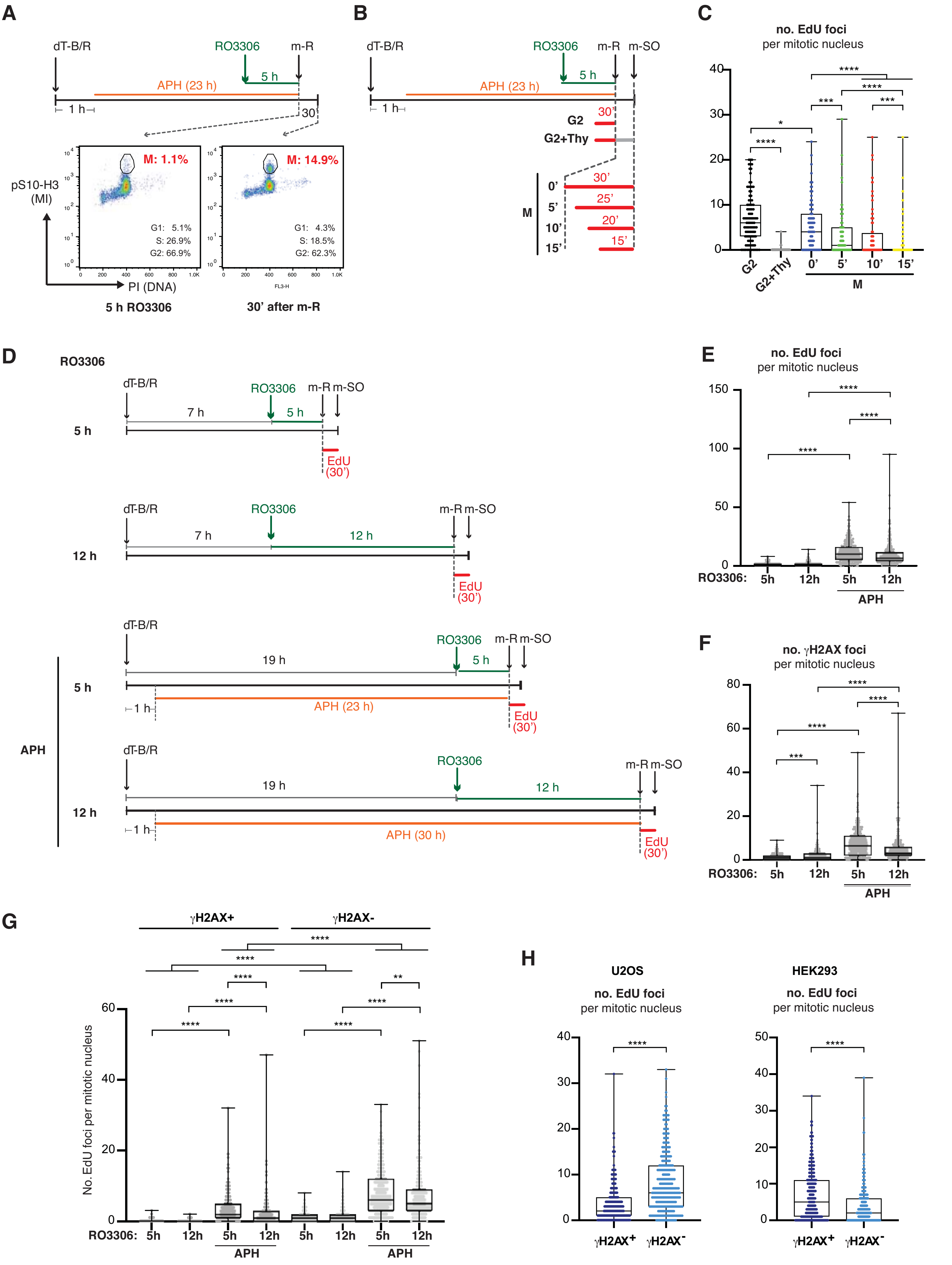

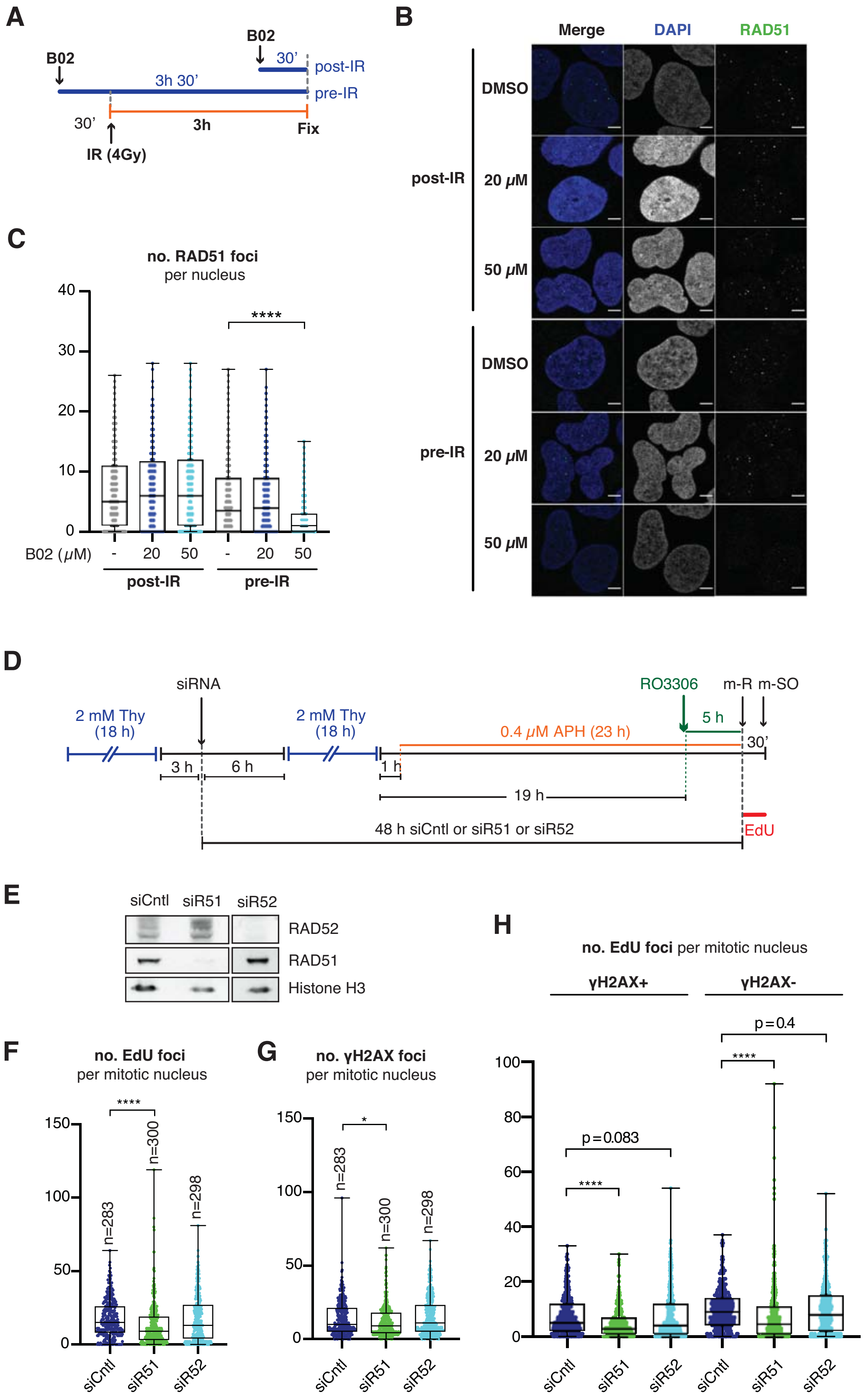
A
B02
B02
30'
3h 30'
post-IR
pre-IR
30'
3h
Fix
IR (4Gy)
B
Merge
DAPI
RAD51
DMSO
post-IR
20 μM
50 μM
DMSO
pre-IR
20 μM
50 μM
C
no. RAD51 foci
per nucleus
****
B02 (μM)
- 20 50 - 20 50
post-IR
pre-IR
D
siRNA
2 mM Thy (18 h)
2 mM Thy (18 h)
RO3306
m-R m-SO
5 h
0.4 μM APH (23 h)
30'
3 h
6 h
1 h
19 h
48 h siCntl or siR51 or siR52
EdU
E
siCntl siR51 siR52
RAD52
RAD51
Histone H3
F
no. EdU foci
per mitotic nucleus
****
n=283
n=300
n=298
siCntl
siR51
siR52
G
no. γH2AX foci
per mitotic nucleus
*
n=283
n=300
n=298
siCntl
siR51
siR52
H
no. EdU foci per mitotic nucleus
γH2AX+
γH2AX-
p=0.4
****
p=0.083
****
siCntl
siR51
siR52
siCntl
siR51
siR52

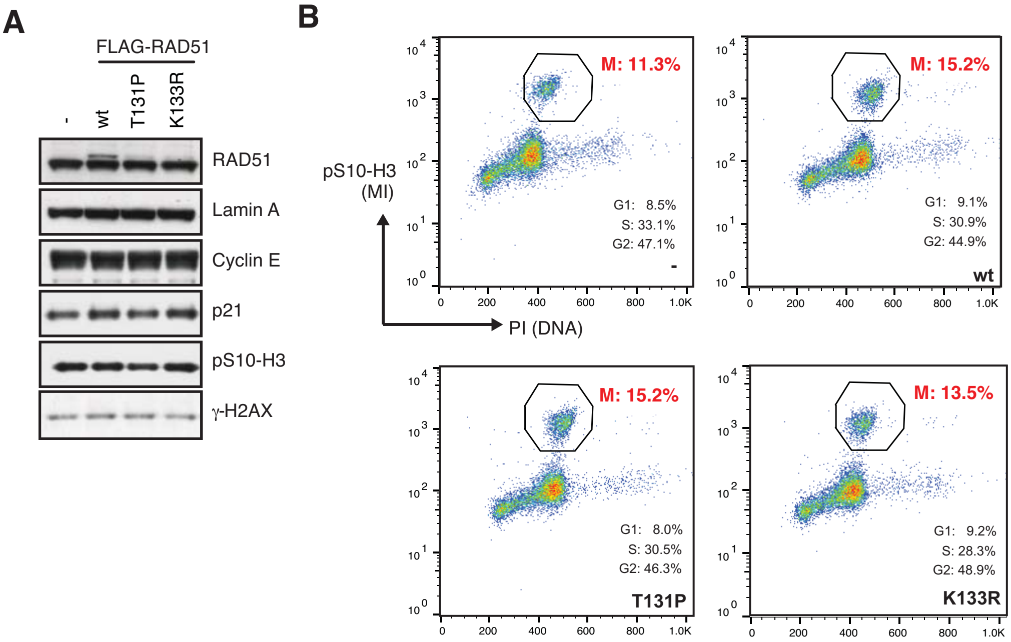
A
FLAG-RAD51
-
wt
T131P
K133R
RAD51
Lamin A
Cyclin E
p21
pS10-H3
γ-H2AX
B
pS10-H3
(MI)
PI (DNA)
M: 11.3%
G1: 8.5%
S: 33.1%
G2: 47.1%
-
M: 15.2%
G1: 9.1%
S: 30.9%
G2: 44.9%
wt
M: 15.2%
G1: 8.0%
S: 30.5%
G2: 46.3%
T131P
M: 13.5%
G1: 9.2%
S: 28.3%
G2: 48.9%
K133R
0
200
400
600
800
1.0K

### Wassing et al., Figure S4

**A**

```
 M   A   M   Q   M   Q   L   E   A   N   A   D   T13 S14 V   E   E   E   S   F   G   P   Q
atg gca atg cag atg cag ctt gaa gca aat gca gat act tca gTG Gaa gaa gaa agc ttt ggc cca caa
```

gRNA

**B**

gRNA-Cas9-GFP + ssDNA repair template
↓
GFP +ve single cell sorting
(96 wells x 10)
↓
PCR screening
↙ ↘
miSeq WB

**C**

| Round | Mutation | Screened | Confirmed |
|---|---|---|---|
| 1 | S14D | 30 | 4 (homozygous) |
| 1 | T13A/S14A | 15 | 0 |
| 2 | T13A/S14A | 19 | 0 |
| 2 | S14A | 27 | 0 |

**D**

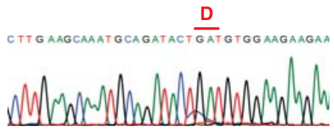


**E**

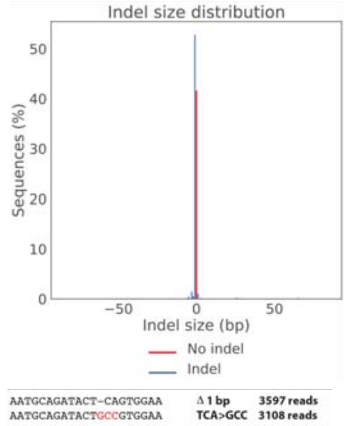


**F**

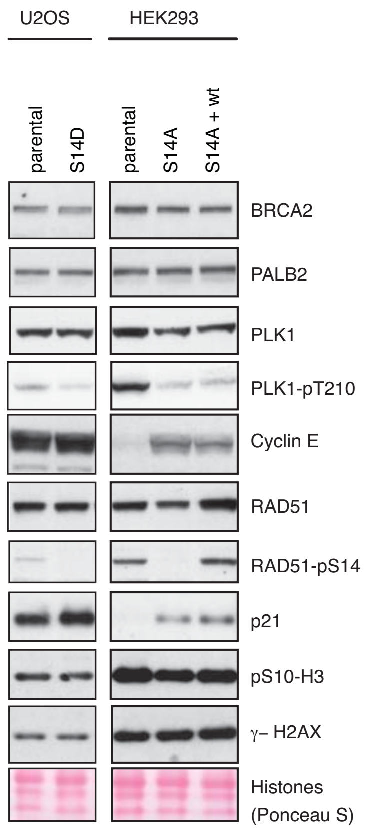


**G**

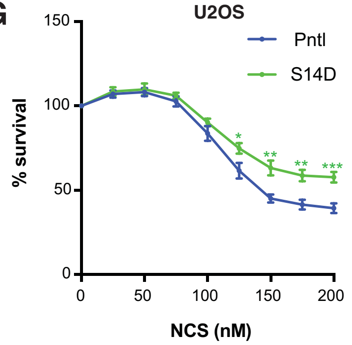


**H**

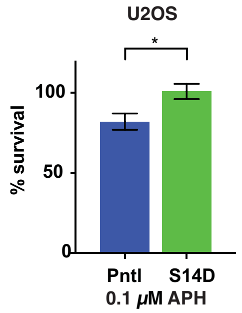


**I**

| U2OS | Division Index NT | Division Index APH (0.5 μM) |
|---|---|---|
| Pntl | 2.40 | 1.06 |
| S14D | 2.37 | 1.28 |

**J**

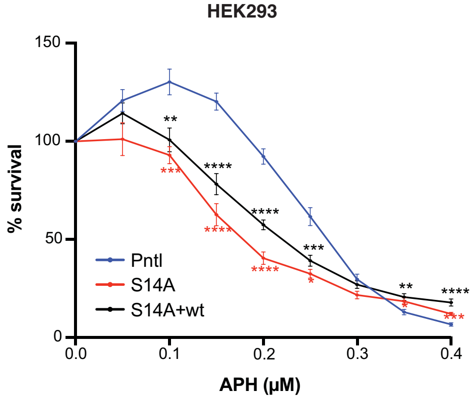


**K**

| HEK293 | Division Index NT | Division Index APH (0.2 μM) |
|---|---|---|
| Pntl | 4.45 | 2.45 |
| S14A | 3.41 | 2.03 |
| S14A +wt | 3.68 | 2.20 |

**L**

S14A vs. S14A+wt

| APH (μM) | p-value |
|---|---|
| 0.15 | 0.062 |
| 0.20 | 0.0007 (***) |
| 0.25 | 0.0799 |
| 0.30 | 0.0705 |
| 0.40 | 0.0094 (**) |

**M**

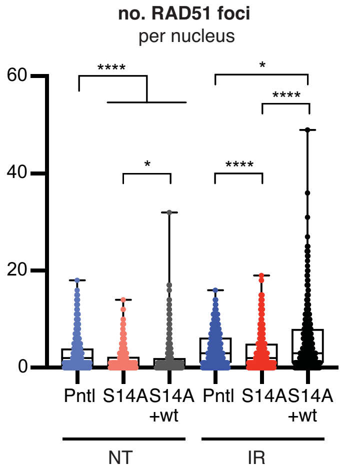


**N**

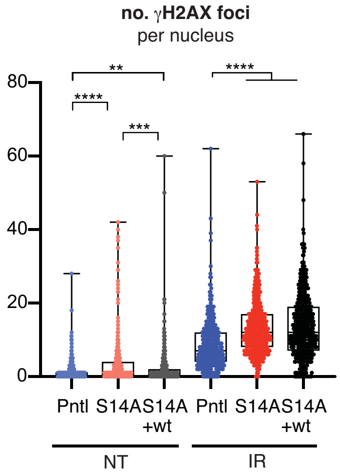

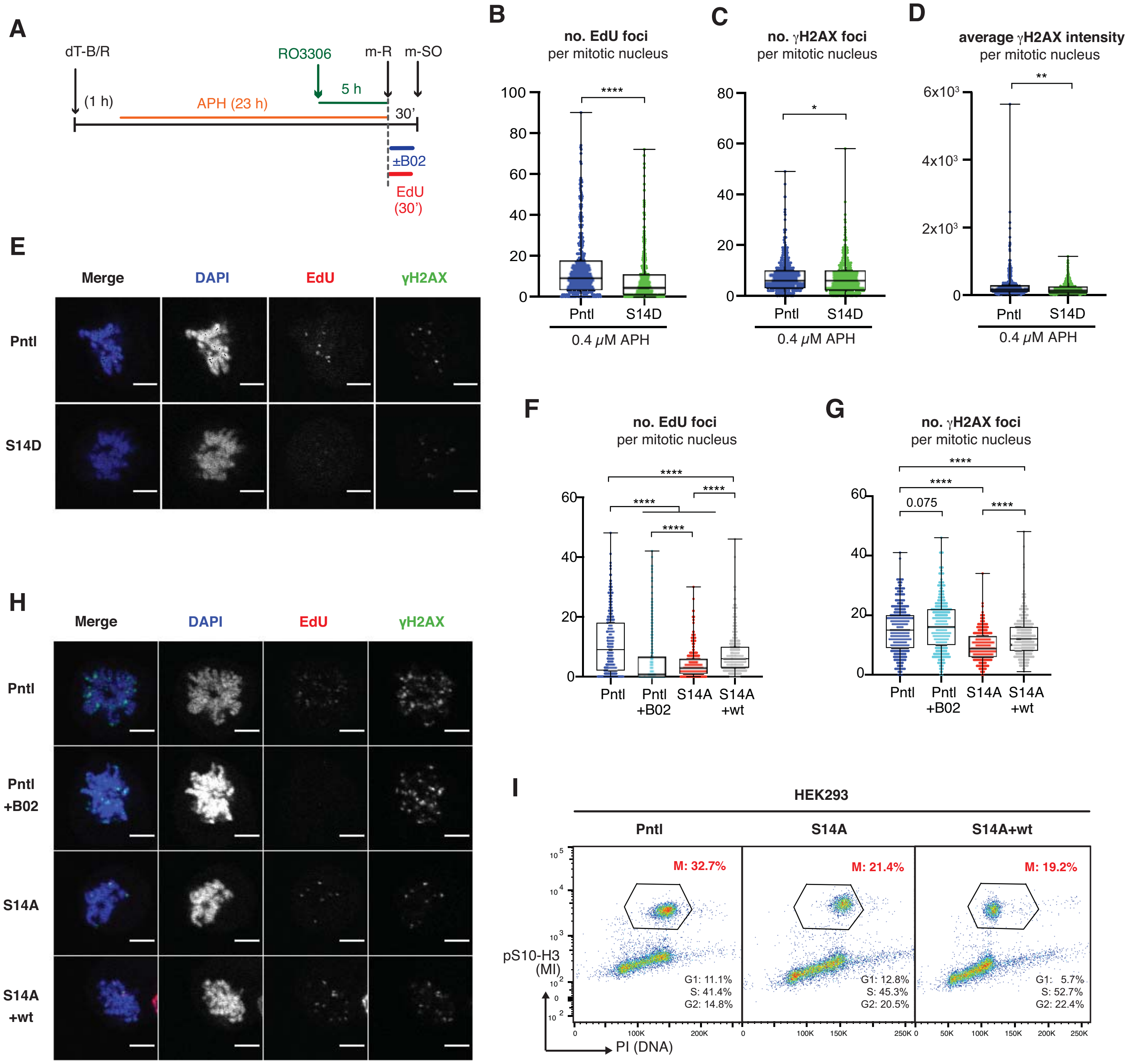
A
dT-B/R
(1 h)
APH (23 h)
RO3306
5 h
m-R
m-SO
30'
±B02
EdU
(30')
B
no. EdU foci
per mitotic nucleus
****
Pntl
S14D
0.4 μM APH
C
no. γH2AX foci
per mitotic nucleus
*
Pntl
S14D
0.4 μM APH
D
average γH2AX intensity
per mitotic nucleus
**
Pntl
S14D
0.4 μM APH
E
Merge
DAPI
EdU
γH2AX
Pntl
S14D
F
no. EdU foci
per mitotic nucleus
****
Pntl
Pntl +B02
S14A
S14A +wt
G
no. γH2AX foci
per mitotic nucleus
****
0.075
Pntl
Pntl +B02
S14A
S14A +wt
H
Merge
DAPI
EdU
γH2AX
Pntl
Pntl +B02
S14A
S14A +wt
I
HEK293
Pntl
S14A
S14A+wt
M: 32.7%
M: 21.4%
M: 19.2%
G1: 11.1%
S: 41.4%
G2: 14.8%
G1: 12.8%
S: 45.3%
G2: 20.5%
G1: 5.7%
S: 52.7%
G2: 22.4%
pS10-H3 (MI)
PI (DNA)

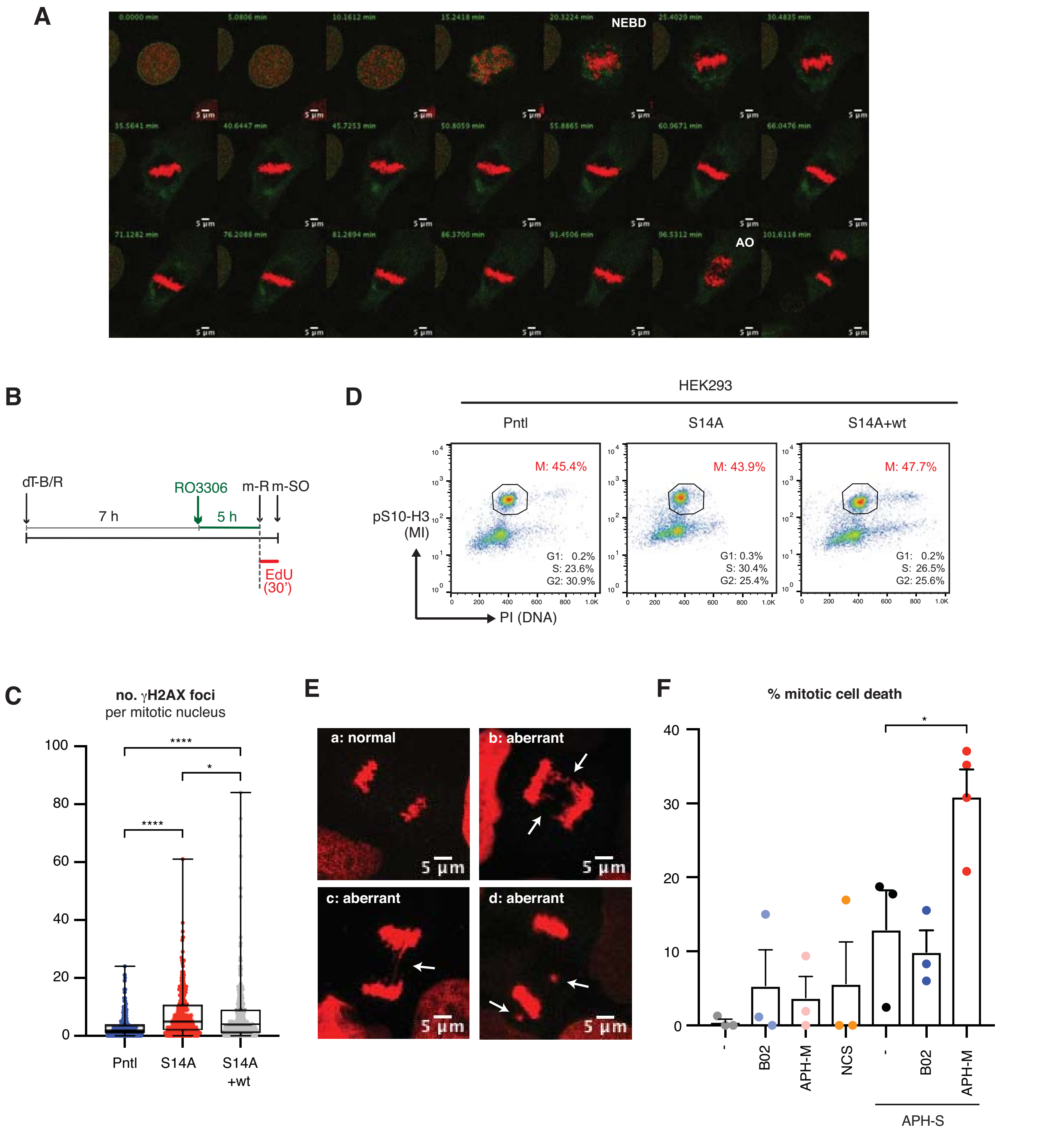
A
NEBD
AO
B
dT-B/R
RO3306
m-R
m-SO
7 h
5 h
EdU (30')
D
HEK293
Pntl
S14A
S14A+wt
M: 45.4%
M: 43.9%
M: 47.7%
pS10-H3 (MI)
PI (DNA)
G1: 0.2% S: 23.6% G2: 30.9%
G1: 0.3% S: 30.4% G2: 25.4%
G1: 0.2% S: 26.5% G2: 25.6%
C
no. γH2AX foci per mitotic nucleus
Pntl
S14A
S14A +wt
E
a: normal
b: aberrant
c: aberrant
d: aberrant
5 µm
F
% mitotic cell death
B02
APH-M
NCS
APH-S

#### Supplemental Figure Legends

**Figure S1: Evaluation of DNA synthesis upon mild replicative stress in mitosis.**

**A.** Schematic diagram of experimental procedure for U2OS cell synchronisation. FACS profiles showing mitotic index (pS10-H3) and DNA content (PI) of cells exposed to RO3306 for 5 hours (left) and at 30 min after mitotic release (m-R) from RO3306 block (right) are also shown. The G1, S and G2 populations were estimated by the Watson Pragmatic algorithm (FlowJo), based on the PI staining of interphase cells.

**B.** Schematic diagram of experimental procedure, as in (A), combined with 30 min EdU-labelling prior to m-R (G2), 30 min EdU-labelling prior to m-R followed by a 30-min 100 µM thymidine chase upon m-R (G2+Thy); 30 min EdU-labelling upon m-R (M, 0'); 25 min EdU-labelling at 5 min upon m-R (M, 5'); 20 min EdU-labelling at 10 min upon m-R (M, 10'); and 15 min EdU-labelling at 15 min upon m-R (M, 15'). Cells were collected 30 min after m-R by mitotic shake-off (m-SO) and analysed for EdU incorporation.

**C.** Quantification of the number of EdU foci per mitotic nucleus following EdU-labelling protocol as in (B). 100 cells were quantified per condition.

**D.** Schematic illustration of cell synchronisation by dT-B/R in the absence or presence of 0.4 µM APH and subsequent incubation in 9 µM RO3306 for 5 h or 12 h. Cells were labelled with EdU for 30 min upon m-R, collected by m-SO and stained for EdU and $\gamma$-H2AX.

**E-F.** Quantifications of the number of EdU foci (E) and $\gamma$-H2AX foci (F) per mitotic nucleus.

**G.** The same data as (E) and (F) is quantified for the number of EdU foci that colocalise with $\gamma$-H2AX ($\gamma$-H2AX$^+$ EdU foci) and the number of EdU foci that do not colocalise with $\gamma$-H2AX ($\gamma$-H2AX$^-$ EdU foci) per mitotic nucleus.

**H.** Mitotic EdU incorporation and $\gamma$-H2AX IF signal was analysed in U2OS and HEK293 cell lines following cell synchronisation in the presence of 0.4 µM APH and 5 hour RO3306-incubation.

Quantifications of the number of EdU foci that colocalise with $\gamma$-H2AX ($\gamma$-H2AX$^+$ EdU foci) and the number of EdU foci that do not colocalise with $\gamma$-H2AX ($\gamma$-H2AX$^-$ EdU foci) per mitotic nucleus are shown. In (E)-(H), data was obtained from 3 independent experiments (100 cells analysed per repeat); n=300 per condition. Data distribution is represented by box-and-whisker plots (whiskers spanning min-max range of values). Mann Whitney test p-values values are shown. Asterisks indicate p value ≤ 0.05 = *; ≤ 0.01 = **; ≤ 0.001 = ***; ≤ 0.0001 = ****.

**Figure S2: Evaluation of RAD51 inhibitor B02 and a comparative analysis between RAD51 and RAD52 depletion for MiDAS.**

**A.** Schematic diagram of experimental procedure to include B02 exposure before (pre-IR) or after (post-IR) irradiation (4 Gy) of asynchronous U2OS cells. 3 hours after irradiation, cells were fixed and stained for RAD51 IF.

**B.** Representative images of cells quantified in (C). Scale bar indicates 5 µm.

**C**. Quantification of the number of RAD51 foci per nucleus. Data were obtained from three independent experiments (100 cells analysed per repeat); n=300 per condition. Data distribution is represented by box-and-whisker plots (whiskers spanning min-max range of values). Mann Whitney test p-values values are shown. Asterisks indicate p value ≤ 0.05 = *; ≤ 0.01 = **; ≤ 0.001 = ***; ≤ 0.0001 = ****.

**D.** Schematic diagram of experimental procedure for U2OS cell synchronisation combined with siRNA transfection, targeting RAD51 (siR51) or RAD52 (siR52), as in Figure 1A.

**E.** Following cell synchronisation according to workflow in (D), the total cell population was collected by trypsinization 30 min after m-R and analysed by Western blot for the depletion of the corresponding protein.

**F-G.** Quantification of the number of EdU foci (F) and $\gamma$-H2AX foci (F) per mitotic nucleus.

**H.** Same data as in (F) and (G), quantified as the number of EdU foci that colocalise with $\gamma$-H2AX ($\gamma$-H2AX$^{+}$ EdU foci) and the number of EdU foci that do not colocalise with $\gamma$-H2AX ($\gamma$-H2AX$^{-}$ EdU foci) per mitotic nucleus. Data were obtained from three independent experiments (approximately 100 cells analysed per repeat); n= the total number of cells quantified per condition as indicated on graph. Data distribution is represented by box-and-whisker plots (whiskers spanning min-max range of values). Mann Whitney test p-values values are shown. Asterisks indicate p value ≤ 0.05 = *; ≤ 0.01 = **; ≤ 0.001 = ***; ≤ 0.0001 = ****.

**Figure S3: Characterisation of U2OS cells expressing RAD51 separation-of-function mutants.**

**A.** Western blot assessing RAD51, Cyclin E, p21, $\gamma$-H2AX, pS10-H3 (mitosis marker) and Lamin A (loading control) levels in asynchronous U2OS Flp-In T-REx cells expressing the indicated FLAG-RAD51 variants from the CMV promoter.

**B.** FACS analysis of mitotic index (pS10-H3) and DNA content (PI) following cell synchronisation according to the experimental procedure depicted in Figure 3A. At 30 min after m-R, the total cell population was collected by trypsinization and analysed by FACS. The G1, S and G2 populations in respective cell lines were estimated by the Watson Pragmatic algorithm (FlowJo), based on the PI staining of interphase cells.

**Figure S4: Evaluation of endogenous RAD51 S14 missense mutant cells.**

**A**. DNA sequence and corresponding amino acid sequence of the RAD51 S14 locus targeted by gRNA for CRISPR/Cas9-mediated gene editing.

**B.** Schematic illustration of workflow for knock-in of RAD51 S14 missense mutations by CRISPR/Cas9 gene editing.

**C.** The total number of CRISPR/Cas9 targeted U2OS clones screened by PCR, and the number of clones validated by Sanger sequencing. Two rounds (Round 1 and 2) of the whole screening procedure, depicted in (B), were attempted.

**D.** Representative Sanger sequencing chromatogram of the RAD51 S14 locus obtained for the U2OS RAD51 S14D homozygous knock-in mutants.

**E.** Graphic representation of the indel size distribution for all reads of the RAD51 S14A locus in the HEK293 knock-in mutant as sequenced by next generation sequencing (miSeq). The corresponding DNA sequences detected are displayed below.

**F.** Asynchronous populations of the U2OS parental cell line, the U2OS S14D knock-in mutant (S14D); the HEK293 parental cell line; the HEK293 S14A knock-in mutant (S14A), as well as the wild-type RAD51 complemented HEK293 S14A knock-in mutant (S14A+wt) were assessed by Western blot against S14 phosphorylated RAD51 (RAD51-pS14), RAD51, BRCA2, PALB2, PLK1, phosphorylated PLK1 (PLK1-pT210), Cyclin E, p21, pS10-H3 (mitotic marker) and $\gamma$-H2AX.

**G.** Cell survival upon exposure to neocarzinostatin (NCS) was assessed by WST-1 assay. Error bars represent SEM; n=9 from three independent clonogenic survival assay experiments (three technical repeats were included per experiment). Significant difference compared to the parental U2OS cell line (Pntl), assessed by unpaired t test, is indicated by asterisks.

**H.** Cell survival upon exposure to 0.1 μM APH was assessed by clonogenic assay. Error bars represent SEM, n=8 from three independent experiments (at least two technical repeats were included per experiment). Significant difference compared to the parental HEK293 cell line (Pntl), assessed by unpaired t test, is indicated by asterisks.

**I.** Proliferation of the CRISPR gene-edited RAD51 S14D mutant and the corresponding U2OS parental cell line (Pntl) was assessed following 96 hour incubation with 0.5 μM aphidicolin (APH) or non-treated control (NT). The division index for each cell line was determined by Cell Trace$^{TM}$ cell proliferation assay based on a single experiment.

**J.** Cell survival of HEK293 cell lines upon exposure to aphidicolin (APH) were assessed by WST-1 assay. Error bars represent SEM; n=9 from three independent experiments (three technical repeats were included per experiment). Significant difference compared to the parental HEK293 cell line (Pntl), assessed by unpaired t test, is indicated by asterisks.

**K.** Proliferation of the CRISPR gene-edited RAD51 S14A mutant, the complemented S14A+wt cell line and the corresponding HEK293 parental cell line (Pntl) was assessed following 96-hour incubation with 0.5 µM aphidicolin (APH) or non-treated control (NT). The division index for each cell line was determined by Cell Trace$^{TM}$ cell proliferation assay based on a single experiment.

**L.** Statistical differences between the HEK293 RAD51 S14A and S14A+wt cell survival, as assayed in (J) are assessed by unpaired t test and summarised.

**M-N.** HEK293 parental (Pntl), S14A and S14A+wt cell lines, at 3 hours after irradiation (IR) at 4 Gy or with no treatment (NT), were assessed for RAD51 foci (J) and $\gamma$-H2AX foci (K). Data were obtained from three independent experiments (150 cells analysed per repeat; n=450 per condition). Data distribution is represented by box-and-whisker plots (whiskers spanning min-max range of values). Asterisks indicate Mann Whitney test p value ≤ 0.05 = *; ≤ 0.01 = **; ≤ 0.001 = ***; ≤ 0.0001 = ****.

**Figure S5: Analysis of MiDAS in CRISPR gene-edited RAD51 S14 knock-in mutants.**

**A.** Schematic diagram of cell synchronisation for the detection of EdU incorporation upon mild replicative stress.

**B-D**. EdU incorporation and $\gamma$-H2AX IF signal were analysed in the parental U2OS cell line (Pntl) and U2OS RAD51 S14D knock-in mutant following cell synchronisation as previously depicted in

(A). Quantification of the number of EdU foci (B) and the number of $\gamma$-H2AX foci per mitotic nucleus (C). Same data as (C), quantified the average $\gamma$-H2AX focus intensity per mitotic nucleus (D). Data were obtained from four independent experiments (100 cells analysed per repeat); n=400 per condition. Data distribution is represented by box-and-whisker plots (whiskers spanning min-max range of values). Asterisks indicate Mann Whitney test p value ≤ 0.05 = *; ≤ 0.01 = **; ≤ 0.001 = ***; ≤ 0.0001 = ****.

**E.** Representative images of the data analysed in (B-D) are shown. Scale bar indicates 5 µm. Pntl indicates the parental U2OS cell line.

**F.** HEK293 cell lines, synchronised as in (A), were released into mitosis in the presence of 10 µM EdU with or without 20 µM RAD51 inhibitor (B02), and mitotic cells collected by m-SO were analysed for EdU incorporation and $\gamma$-H2AX signal. Pntl indicates the parental HEK293 cell line. Representative images are shown, scale bar indicates 5 µm.

**G-H.** Quantification of the number of EdU foci (G) and $\gamma$-H2AX foci (H) per mitotic nucleus. Pntl indicates the parental HEK293 cell line. Data were obtained from three independent experiments (100 cells analysed per repeat); n=300 per condition. Data distribution is represented by box-and-whisker plots (whiskers spanning min-max range of values). Asterisks indicate Mann Whitney test p value ≤ 0.05 = *; ≤ 0.01 = **; ≤ 0.001 = ***; ≤ 0.0001 = ****.

**I.** HEK293 parental (Pntl), S14A and S14A+wt cell lines, synchronised as in (A). The total cell population was analysed by FACS for mitotic index (pS10-H3) and DNA content (PI). The G1, S and G2 populations in respective cell lines were estimated by the Watson Pragmatic algorithm (FlowJo), based on the PI staining of interphase cells.

**Figure S6: Impact of RAD51 and MiDAS on mitotic progression.**

**A.** Representative image of mitotic progression analysed in U2OS cells following cell synchronisation as in Figure 5F. Frames displaying nuclear envelope breakdown (NEBD) and anaphase onset (AO) are indicated.

**B.** Schematic diagram of cell synchronisation for indicated HEK293 cell lines.

**C.** Quantification of the number of $\gamma$-H2AX foci per mitotic nucleus, detected in mitotic cells collected by m-SO. Pntl indicates the parental HEK293 cell line. Data were obtained from three independent experiments (100 cells analysed per repeat); n=300 per cell line. Asterisks indicate Mann Whitney test p-value ≤ 0.05 = *; ≤ 0.01 = **; ≤ 0.001 = ***; ≤ 0.0001 = ****..

**D.** The total cell population at 30 min after m-R was collected by trypsinization and analysed by FACS for mitotic index (pS10-H3) and DNA content (PI). The G1, S and G2 populations in respective cell lines were estimated by the Watson Pragmatic algorithm (FlowJo) based on the PI staining of interphase cells.

**E.** Representative images of normal (a) and aberrant mitotic chromatin (b-d) as quantified in Figure 5G. Arrows indicate lagging chromosomes, chromatin bridges or micronuclei.

**F.** The experimental data, analysed for Figure 5G, was evaluated for the percentage of cells that die before completing mitosis. Asterisks indicate unpaired t test p value ≤ 0.05 = *; ≤ 0.01 = **; ≤ 0.001 = ***; ≤ 0.0001 = ****.

#### Supplemental Movie Legends

**Supplemental Movie 1: Representative mitotic progression of HEK293 cell.** Following synchronisation as depicted in Figure 5D.

**Supplemental Movie 2: Representative mitotic progression of HEK293 S14A cell.** Following synchronisation as depicted in Figure 5D.

**Supplemental Movie 3: Representative mitotic progression of HEK293 S14A+wt cell.** Following synchronisation as depicted in Figure 5D.

**Supplemental Movie 4: Representative mitotic progression of U2OS cell.** Following synchronisation in the absence of aphidicolin as depicted in Figure 5F.

**Supplemental Movie 5: Representative mitotic progression of U2OS cell.** Following synchronisation in the presence of aphidicolin as depicted in Figure 5F

**Supplemental Movie 6: Representative mitotic progression of U2OS cell in the presence of 20 µM B02.** Following synchronisation in the absence of aphidicolin as depicted in Figure 5F.

**Supplemental Movie 7: Representative mitotic progression of U2OS cell in the presence of 20 µM B02.** Following synchronisation in the presence of aphidicolin as depicted in Figure 5F

**Supplemental Movie 8: Representative mitotic progression of U2OS cell in the presence of 2 µM aphidicolin.** Following synchronisation in the absence of aphidicolin as depicted in Figure 5F.

**Supplemental Movie 9: Representative mitotic progression of U2OS cell in the presence of 2 µM aphidicolin.** Following synchronisation in the presence of aphidicolin as depicted in Figure 5F

**Supplemental Movie 10: Representative mitotic progression of U2OS cell in the presence of 2 µM AZ3146.** Following synchronisation in the absence of aphidicolin as depicted in Figure 5F.

**Supplemental Movie 11: Representative mitotic progression of U2OS cell in the presence of 2 µM AZ3146.** Following synchronisation in the absence of aphidicolin as depicted in Figure 5F.

**Supplemental Movie 12: Representative mitotic progression of U2OS cell in the presence of 50 nM neocarzinostatin.** Following synchronisation in the absence of aphidicolin as depicted in Figure 5F

#### Supplemental Methods

##### Table S1

| ssDNA repair template | |
|---|---|
| S14D | CCAAAATTAGCAACTAACCACATACCTCTAACCGTGAAATGGGTTGTGGCCAAAGCTT TCTTCTTCCAC**ATC**AGTATCTGCATTTGCTTCAAGCTGCATCTGCATTGCCATTACTGAA AAATACAAATGCTTATCAGTATAAACACTAG |
| S14A | CCAAAATTAGCAACTAACCACATACCTCTAACCGTGAAATGGGTTGTGGCCAAAGCTT TCTTCTTCCAC**GGC**AGTATCTGCATTTGCTTCAAGCTGCATCTGCATTGCCATTACTGAA AAATACAAATGCTTATCAGTATAAACACTAG |

##### CRISPR/Cas9-mediated editing of *RAD51* gene

Cells were co-transfected with (i) a plasmid carrying Cas9, GFP and a guide RNA targeting RAD51 S14 (AGCAAATGCAGATACTTCAGTGG) and (ii) the relevant ssDNA template (**Table S1**), using Lipofectamin LTX with Plus Reagent (ThermoFisher Scientific) according to manufacturer's instructions. At 24 hours after transfection, GFP-expressing cells were sorted by FACS (MoFlo XDP, Beckman Coulter). Single clones were expanded and genomic DNA was extracted for PCR amplification of the targeted RAD51 S14 locus. Clones were screened for the desired mutation using forward primers that are perfectly complementary to either the wildtype S14 locus (CTTGAAGCAAATGCAGATACTTCA), or the mutated S14D (CTTGAAGCAAATGCAGATACTGAT) or S14A (CTTGAAGCAAATGCAGATACT-GCC) locus. Clones that generated a convincing PCR product with the mutation-specific primer, but not the wild-type primer, were further verified for the knock-in mutation by Sanger sequencing. For heterozygous clones, next-generation sequencing (Illumina MiSeq Next Generation Sequencer) was used to determine the allele-specific sequences at the RAD51 S14 locus. Verified clones were further tested for the absence of RAD51 phosphorylation by Western Blot.

**Antibodies**

Primary antibodies used for Western blotting were: anti-phospho-Histone H2A.X (Ser139), clone JBW301 (1:1000, Merck Millipore, 05-636); anti-Histone H3 (1:1000, Bethyl Laboratories, A300-822A); anti-phospho-Histone H3 (Ser 10) (1:1000, Merck Millipore, 06-570); anti-PLK1 (1:1000, Bethyl Laboratories, A300-251A); anti-phospho-PLK1 (T210) (1:1000, BD Biosciences, 558400); anti-BRCA2 (1:1000, Sigma-Aldrich, OP95); anti-PALB2 (1:1000, Biorbyt, orb412704); anti-cyclin E (1:1000, Santa Cruz Biotechnology, HE12); anti-phospho-RAD51 S14 (1:1000, generated as described in Yata et al. 2012), anti-p21 (1:1000, Cell Signalling, 12D1), anti-MUS81 (1:500, Santa Cruz, sc-47692), anti-RAD52 (1:200, Santa Cruz, sc-365341), anti-RAD51 (7946, generated as described in Yata et al. 2014). Primary antibodies used for IF analysis were: anti-phospho-Histone H2A.X (Ser139), clone JBW301 (1:1000, Merck Millipore, 05-636); anti-RAD51(1:1000, generated as described in Yata et al. 2014); anti-RPA (1:5000, Bethyl Laboratories, A300-244A). Anti-phospho-Histone H3 (Ser 10) (1:100, Merck Millipore, 06-570) was used to analyse mitotic index by flow cytometry.

**Cell Trace$^{TM}$ Cell proliferation assay**

U2OS cells were stained following the manufacturer's protocol (Cell Trace$^{TM}$ Cell Proliferation kit, ThermoFisher Scientific). 4x$10^6$ cells were incubated in 0.5 µM Cell Trace Far red solution at a cell density of $10^6$/ml at 37 $^oC$ for 20 min in PBS. 4 volumes of complete medium were added, and cells were incubated for 5 min at 37 $^oC$ to absorb unbound dye. Unbound dye was removed, and cells were seeded in fresh complete medium at a density of 300 000 cells for U2OS and 500 000 cells for HEK293 per 10-cm dish. Cells were incubated in aphidicolin at the indicated concentrations or left untreated. Untreated cells were harvested by trypsinization 24h after seeding to determine the undivided population. Untreated and aphidicolin-treated samples were harvested by trypsinization

96h after seeding to assess the effect of aphidicolin on proliferation. Cells were stained with DAPI (50 ng/ml) to distinguish live and dead cells. Cells were analysed using a Cytek DxP8 Flow cytometer. The obtained proliferation curve was fitted using FlowJo analysis software (version 10.5.3). Parameters were fixed at 0.5 Peak Ratio and Peak was defined CV based on the undivided population as analysed at 24h.

**CO-FISH**

Metaphase spreads were rehydrated in PBS before being treated with 0.5 mg/ml RNAse A (Sigma-Aldrich) in PBS for 10 min at 37 °C. The spreads were next stained with 0.5 µg/ml Hoechst 33258 (Sigma-Aldrich) in 2x SSC for 15 min at room temperature and exposed to 6500 J/m$^2$ 365 nm UV light. To degrade the BrdU:BrdC-containing DNA strands, the metaphase spreads were incubated with 10 U/µl Exonuclease III (Promega) in the manufacturer's buffer at 37 °C for 30 min. The spreads were dehydrated by consecutive incubations in 70%, 90% and 100% ethanol and air-dried. The spreads were next incubated with the PNA probe (F3004, PNABio) in hybridising solution (10 mM Tris-HCl pH 7.2, 70% formamide and 0.5% w/v blocking reagent (Roche)) for 2 hours. The metaphase spreads were briefly rinsed in hybridisation wash 1 (10 mM Tris-HCl pH7.2, 70% formamide and 0.1% BSA) and incubated with the complementary PNA probe (F3009, PNABio) in hybridising solution. The metaphase spreads were washed in hybridisation wash 1 for 15 min, followed by three 5 min washes in hybridisation wash 2 (0.1 M Tris-HCl pH 7.2, 0.15 M NaCl, 0.08% Tween-20). DNA was counterstained by adding 0.7 µg/ml DAPI (Sigma Aldrich) to the second wash. The spreads were dehydrated by consecutive incubations in 70%, 90% and 100% ethanol and air-dried, then finally mounted in ProLong Gold antifade reagent (Life Technologies).
